# LD Matrix Approximations for Scalable Analysis of High-dimensional Genetic Data

**DOI:** 10.1101/2025.09.16.676478

**Authors:** Ulises Bercovich, Shadi Zabad, Simon Gravel

**Affiliations:** Department of Mathematical Sciences, University of Copenhagen; School of Computer Science, McGill University; Department of Human Genetics, McGill University

## Abstract

Linkage disequilibrium (LD) matrices are an essential part of many statistical genetics methods. However, their high dimensionality makes their computation and storage impractical for large genomic data. Common sparse approximations, such as banded matrices, come at the expense of losing the positive semi-definite (PSD) property, a critical quality that ensures numerical stability of many downstream analyses. Conversely, methods that guarantee a PSD approximation, like block-diagonal approaches, require coarse approximations of the LD structure. In this work, we present a novel method to approximate an LD matrix with a sparse, banded matrix that is guaranteed to be PSD while preserving the correlation structure within the band. This is done via a reformulation of the nearest correlation matrix problem using the Cholesky decomposition, which implicitly imposes the PSD property in a highly scalable parallel approach. On whole-chromosome data from the 1000 Genomes Project and the UK Biobank, our method builds sparse positive semi-definiteness that are more more accurate than either block-diagonal or shrinkage estimators.

## Introduction

Linkage disequilibrium (LD), the non-random association of alleles at pairs of genetic loci, is a fundamental quantity in the analysis of genomic data [1–3]. It is influenced by a variety of biological and evolutionary processes such as recombination, selection, and population structure, among others [2–4]. For this reason, LD itself is often used to infer the characteristics of a population, such as their history and admixture events [5–7].

For medical and statistical genetics, LD presents both opportunities and challenges. In addition to paving the way for the early gene-mapping studies, LD was instrumental in laying the foundations for array-based Genome-wide Association Studies (GWASs) [8–10]. On the other hand, in recent years, with the increasing availability of dense genomic datasets [11, 12], this lack of independence between loci complicates downstream analyses. For instance, non-causal genetic variants often show association with a particular disease due to linkage with nearby causal variants. Statistical fine-mapping methods account for LD to narrow down the list of potential causal variants, but reduction to a single candidate is often impossible [13]. In addition, heterogeneous and long-range LD patterns can bias the inference of variant effect sizes for a given phenotype, which can degrade the accuracy and portability of polygenic scores in practical settings [14–16].

Thus, LD matrices, which record the pairwise correlations between a set of genetic variants, play a major role in many modern statistical genetics methods [13, 17–19], especially those that are based on GWAS summary statistics [20]. To see why, it helps to review a commonly-used linear model and associated statistical quantities. For a sample of *n* individuals with phenotype measurements ***y*** ∈ ℝ^*n*^ and corresponding genotype data for *m* variants **G** ∈ ℝ^*n*×*m*^, the linear model is defined as:

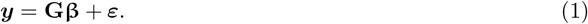

Here, **β** is a *m* × 1 vector that records the effect size of each variant on the phenotype and ***ε***, of dimension *n* × 1, captures the residual effects. If we assume that the genotype matrix and the phenotype vector are standardized column-wise to have zero mean and unit variance, a common measure of the in-sample LD is the *m* × *m* matrix with pairwise Pearson correlation coefficients, defined as 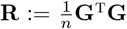. For given set of effect sizes **β**, we also define the Mean Squared Error (MSE), which corresponds to the mean of the squared residuals in the sample,

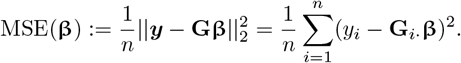

This quantity figures centrally in many statistical analyses of genetic data. Many of the methods discussed above aim to identify effect sizes *β* that best explain the phenotype or, equivalently, that minimize the MSE or an objective function that includes the MSE.

The MSE can be evaluated without access to individual-level data if we have access to the marginal GWAS effect sizes 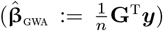 and the in-sample LD matrix **R**, making it possible to provide rich genetic analysis relying on these two summary statistics:

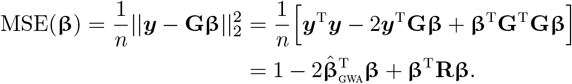

While this is useful theoretically, working with the full in-sample LD matrix in modern genetic analyses is computationally expensive, especially for genome- or chromosome-wide inference methods where data for tens of thousands or millions of genetic markers are processed jointly [15, 16]. For storage and memory, the cost is roughly quadratic in the number of variants 𝒪 (*m*^2^), and for some downstream inference tasks, such as eigen-decomposition, the computational time scales cubically 𝒪 (*m*^3^) with the number of variants.

To get around these computational challenges, researchers have developed a number of different approximations to the in-sample LD matrix. The first major class of methods constructs sparse, banded, or block-diagonal versions of the in-sample LD matrix (Figure 1) [21–25]. Others pursued a shrinkage-based estimator of the LD matrix, which can also produce sparse estimates [19, 26]. These two classes of methods leverage the common understanding that LD tends to decay with genomic distance [1]. More recently, some researchers proposed representing relatively-independent sub-blocks of the LD matrix with low-rank approximations [15, 27]. Others have shown that many of the sparsification strategies can be combined with data compression techniques, such as quantization, to significantly reduce the storage and memory requirements of large-scale LD matrices [16, 28, 29]. These latest approaches enabled scaling statistical genetics methods from 1-2 million to over a dozen million genetic variants [15, 24, 25].

**Figure 1.**
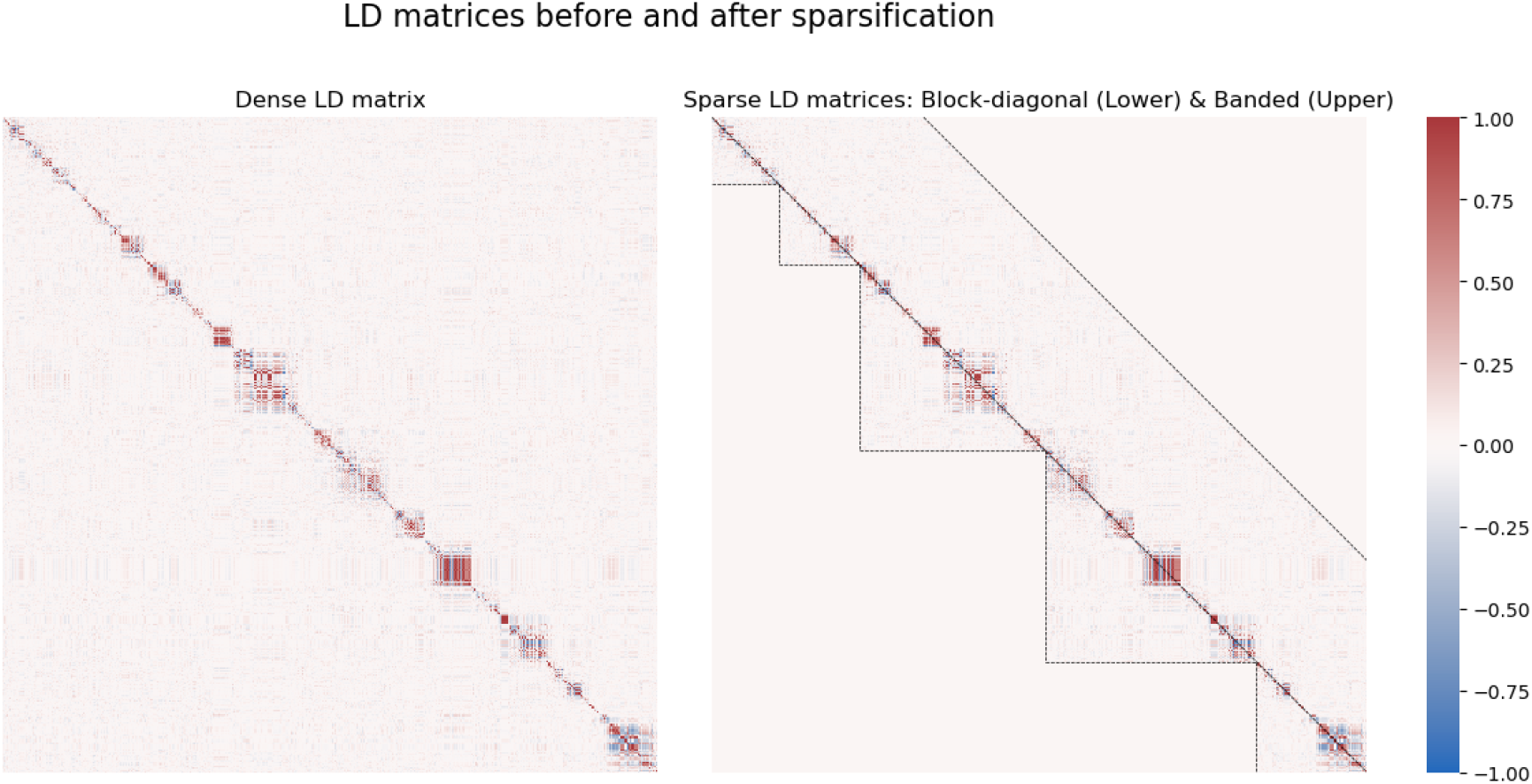
Comparison of a dense Linkage Disequilibrium (LD) matrix with two common sparsification methods on a region of Chromosome 22. The left panel displays the full, dense LD matrix, where each pixel represents the correlation coefficient (*r*) between a pair of SNPs. The right panel shows two methods for creating a sparse matrix to improve computational efficiency. The lower triangle shows a block-diagonal structure, which preserves LD information only within identified haplotype blocks (demarcated by dashed lines). The upper triangle shows a banded structure, which retains LD information only for SNPs within a fixed distance of each other.

Despite these successes, the approximations come with their own set of limitations. For example, block-diagonal LD matrices ignore non-negligible correlations around the boundaries of the blocks, which can complicate resolving genetic association signals in these regions. Banded LD approximations, on the other hand, often result in non-positive semi-definite matrices [16], a critical property that must be preserved for stable and accurate numerical algorithms [16, 27, 30]. A matrix is said to be positive semi-definite (PSD) if it is symmetric (**R** = **R**^t^) and satisfies the inequality,

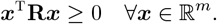

Recent work has shown that many common approaches for estimating or approximating LD lead to non-PSD matrices, including quantization (i.e. aggressive compression) or ignoring missing values in the original genotype matrix [16]. Other approaches, such as shrinkage, may be difficult to scale to modern genomic datasets.

In applied statistics and linear algebra, the problem of approximating matrices while retaining desirable numerical properties, such as sparsity or positive semi-definiteness, has been studied extensively [30–34]. In the “Nearest correlation matrix” literature, the problem is framed as a constrained high-dimensional optimization problem where the goal is to minimize the following objective [30, 33]:

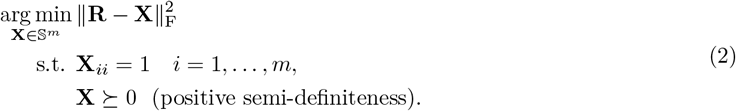

where **R** is a target matrix, 𝕊^*m*^ is the space of symmetric matrices of size *m* × *m*, and ∥ · ∥ _F_ is the Frobenius norm. One major advantage of this formulation is that the objective is convex, which implies the existence of a unique solution. Nonetheless, many of the proposed algorithms, such as the alternating projections method of Higham et al. [30], are not scalable beyond several thousand variants, due to the iterated computation of the eigen-decomposition.

In this work, we present a new method to solve the nearest correlation matrix problem with a banded structure using a reparameterization of the problem similar to the approaches presented in [31, 32, 35]. To avoid expensive eigen-decomposition, we use gradient descent to find matrices that are close to our target matrix and that admit a Cholesky decomposition [36]. Each step of the algorithm improves agreement with the original LD matrix while preserving sparsity and positive semi-definiteness. Furthermore, the algorithm can be initialized using the Cholesky decomposition of the commonly-used block-diagonal approximation of the LD matrix, guaranteeing no deterioration relative to the state of the art. To illustrate the numerical properties and scalability of the proposed algorithm, we applied it to real LD matrices derived from 1000 Genomes Project data [37] and showed that it provides accurate and numerically stable estimates.

## Methods

### Overview of the proposed method for approximating in-sample LD

Given an *n* × *m* genotype matrix **G** of *n* individuals with *m* genetic variants, let **R** be the *m* × *m* in-sample LD matrix, with **R**_*ij*_ = Cor(**G**_·*i*_, **G**_·*j*_) being the sample Pearson correlation coefficient between pairs of variants *i* and *j*. Let **R**_*b*_ be the banded form of **R**, which is the matrix given by

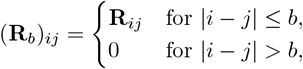

where *b* is the bandwidth of **R**_*b*_. While the original matrix **R** is positive semi-definite (PSD) by construction, **R**_*b*_ might not inherit the PSD property. To correct for this, following earlier work [27], we formulate the following optimization problem

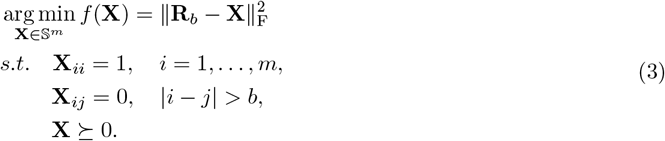

This is a particular case of the nearest correlation matrix problem [30] with the added constraint that the matrix retains a banded and sparse structure. Since the solution **X**^∗^ to Problem (3) is PSD, it is guaranteed to have a Cholesky-like decomposition. As we show in the supplement (Lemma S2), the problem can therefore be reformulated as:

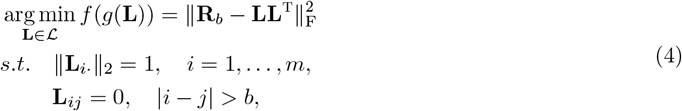

where ℒ is the space of lower triangular matrices and *g*(**L**) := **LL**^T^. The feasible set ℱ_*Chol*_ for this problem is the intersection ℒ_*b*_ ∩ 𝒮^1^, where ℒ_*b*_ := {**L** ∈ ℒ | **L**_*ij*_ = 0, |*i* − *j*| > *b*} is the space of banded lower triangular matrices and 𝒮^1^ := {**M** ∈ ℝ^*m*×*m*^ | ∥**M**_*i*·_∥_2_ = 1} is the Riemannian manifold of real matrices with rows of unit norm. Thus, the feasible set ℱ_*Chol*_ is also a Riemannian manifold.

An advantage of Problem (4) relative to Problem (3) is that we do not have to verify the PSD condition. Many existing methods repeatedly compute the nearest PSD matrix, which involves an expensive computation of the eigen-decomposition. A downside of Problem (4) is that we lose the convexity of the objective function *f* (*g*(**L**)). For example, for a matrix **L**, its negative given by − **L** has the same objective function, but **0**, the midpoint between the two, has not. Non-convexity makes it challenging to identify the global optimum of the objective function. However, gradient-based approaches can still improve the objective from most starting points.

To solve Problem (4), we therefore consider a gradient descent method. The gradient of the objective function is

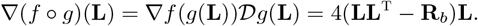

Because the feasible set ℱ_*Chol*_ is a Riemannian manifold, taking steps from a feasible solution along the gradient is not guaranteed to stay within ℱ_*Chol*_. Gradient descent on Riemannian manifolds can be implemented by computing the gradient on the tangent space of manifold, taking a step, and projecting back to the manifold [38]. We therefore need to define projection operations 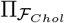 to the feasible set. Since ℱ_*Chol*_ = ℒ_*b*_ ∩ 𝒮^1^, we separately compute the projections 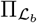 and 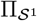 to each set, and use that, in this particular case,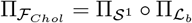.

The projection 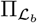 into ℒ_*b*_ is easily computed by keeping the elements on the lower triangular part within the band and inserting zeros outside the band on the lower triangular part of the matrix and on the strictly upper triangular part. The set 𝒮^1^ includes matrices **L** where each row is on the *m*−dimensional sphere of radius 1. Therefore, to project the gradient into tangent space of the feasible manifold, we have to do a row-wise subtraction of the normal component to the sphere, which is done by computing

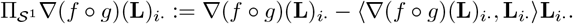

Based on the computed projections and the known gradient, we can define the gradient on the Riemannian manifold

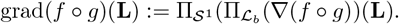

The Riemannian gradient grad(*f* ° *g*)(**L**) gives a direction of maximum increase of the objective function at point **L** within the manifold. However, taking a finite step along the gradient does not guarantee that we stay within the manifold. As typical in Riemannian gradient descent, we take finite steps along the Riemannian gradient followed by projection into the feasible space. This algorithm can be found in Algorithm 1. The algorithm returns a banded matrix **L**^∗^ that is used to calculate a feasible solution for Problem 3 by using the map *g*(**L**^∗^).

#### Algorithm 1

Cholesky-based nearest banded correlation matrix

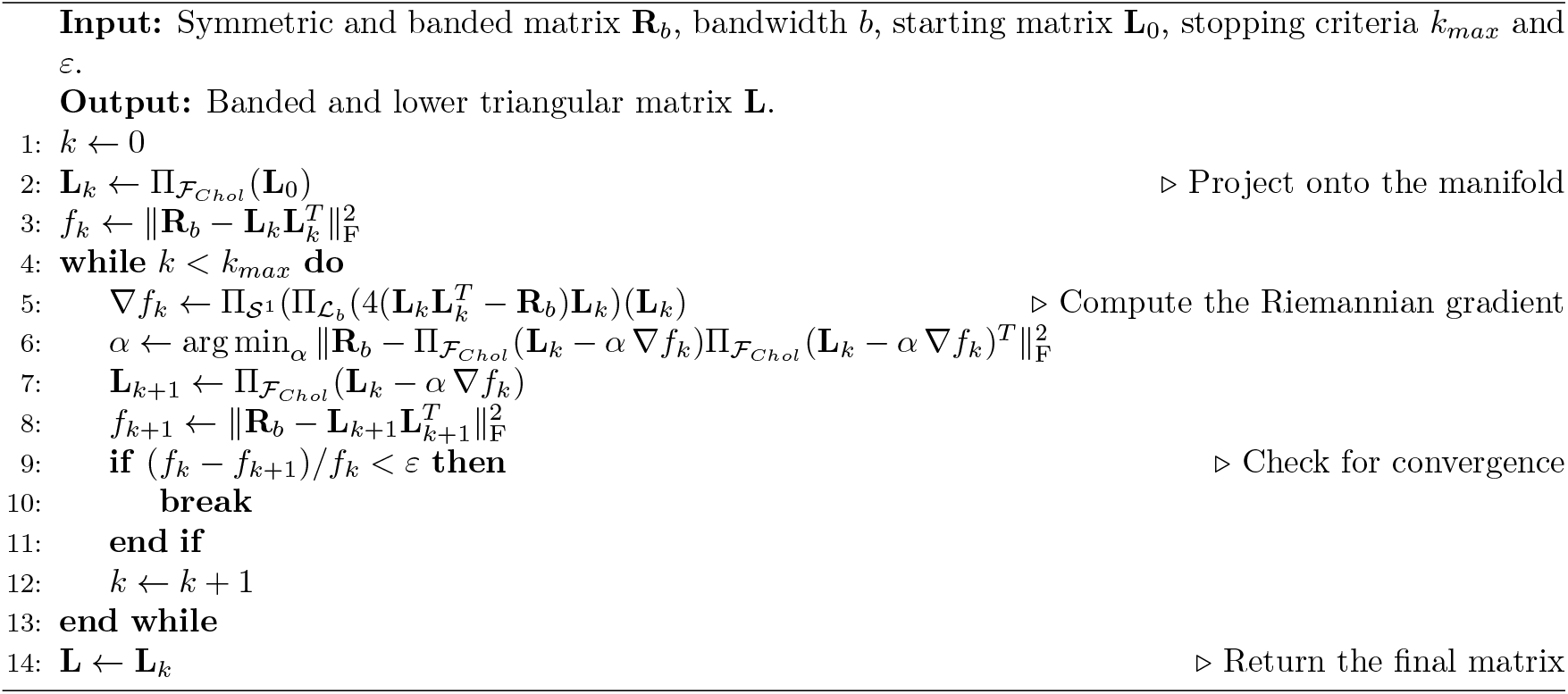

This procedure can use previously determined PSD approximations of the matrix **R** as a starting point. Popular examples are based on extracting a block-diagonal matrix 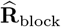 from **R** [21, 39], which is based on the fact that every block-diagonal mask of a PSD matrix is also PSD. Afterwards, we can calculate the Cholesky decomposition of 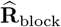 and use it as the starting point for the algorithm. This ensures that the gradient descent method improves upon the block-diagonal PSD approximation of **R**_*b*_. In the absence of a starting point, the identity matrix or a shift of **R**_*b*_ based on its most negative eigenvalue can be used as a feasible starting solution.

In contrast to algorithms that require the eigen-decomposition, the execution time of each iteration of our method is highly reduced by avoiding power methods. The computational cost of each iteration can be explicitly calculated by looking at the most expensive step: the computation of 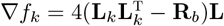. As the matrices are banded and the result of this multiplication is projected to the space of lower triangular matrices with bandwidth *b*, we only have to compute approximately *m* × *b/*2 elements. This results in an iteration computational cost of 𝒪 (*mb*^2^). Computation of the step size *α* is also 𝒪 (*mb*^2^) as long as we use methods that avoid the computation of the Hessian matrix.

While we have presented an implementation for a banded matrix, the approach can be generalized to account for variable bandwidth, for instance by considering changes in the recombination rate or SNP density. This is done by relaxing the projection given by 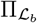 and allowing projecting onto a less rigid space. For example, ℒ_*b*_ can be defined using a variable bandwidth given by a local recombination rate or the convex hull of a block-diagonal mask, as presented in Supplementary Figures S6 and S7. This is based on the fact that Cholesky decompositions also preserve the sparsity structure of matrices with sparse envelopes (Lemma 4.2.1 in [40]).

## Methods to compute step size

### Backtracking line search

For the matrices where the LD matrices can be stored in RAM, we studied a backtracking line search with the Armijo-Goldstein condition as the stopping criteria [41, 42]. To improve the speed of the search, we use an adaptive initial step size. This is done via a warm start, where on the first iteration, we begin with an initial step size of *α*_0_. For the following iterations, we use the previous accepted step size *α*_*k*−1_ as a warm start.

### Parallel Adam optimizer

On large chromosomes, the dimensionality of the matrices **R**_*b*_ and **L** can be a burden when computing 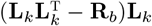. Given the banded form of the matrices, where we assume that *b* << *m*, we can compute the gradient on a block without the need to access the full matrices. To do this, we use the Adam optimizer [43] where the step size is given by quantities that can be computed in parallel on each block. This allows us to avoid handling the full matrix and only use, for each block, the information within the block and its adjacent blocks.

A detailed description of these approaches can be found in the Supplementary material on Algorithms S1 and S2. A speed and convergence comparison is presented in the Supplementary Section S2.

## Genotype data and linkage-disequilibrium approximations

To demonstrate the properties and advantages of the proposed algorithm for LD approximation, we conducted various experiments using real genotype data from The 1000 Genomes Project [37]. Specifically, we used genotype data for a subset of individuals of European ancestry (*n* = 378) along with a restricted subset of HapMap3 variants on Chromosome 22, totaling *m* = 15, 938 Single Nucleotide Polymorphisms (SNPs). This dataset is available for download and small-scale experimentation via the `magenpy` software package [24], which comes with utilities for reading the genotype data and converting it to dense `numpy` [44] or sparse `scipy` matrices [45]. To demonstrate the scalability of the method, we used pre-computed LD matrices of the 22 human autosomes from [16], downloaded directly from its public repository. These data were originally derived from individuals of European ancestry in the UK Biobank [46].

Computing sparse LD matrices with banded, block-diagonal, or shrinkage approximations was done with routines provided by the `magenpy` software package [24]. In its default setting, `magenpy` computes LD by standardizing the genotype matrix **G** column-wise and then computing the in-sample LD matrix, as defined previously, with 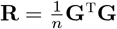. Once the in-sample LD matrix is constructed, then the approximation routines are applied to sparsify or shrink the matrix, with parameters defined by the user. For the block-diagonal approximation, we used LD blocks defined by `LDetect` [21]. For banded LD matrices, we used a denser distance based on the largest block from the previous method `LDetect` to delineate a fixed bandwidth, and a sparser adaptive band based on centiMorgans (cM) to have the same quantity of non-zero elements as the block-diagonal approach (more details in Table S1). For the shrinkage estimator, we specified the sample size for the genetic map as *N*_*e*_ = 183, the effective population size as *N*_*e*_ = 11400, and the default shrinkage cutoff as 10^−3^.

## Metrics used for comparing LD approximations

To quantify the accuracy of each LD approximation algorithm, we started with a normalized version of the objective function defined in Equation (2) and computed the relative Frobenius error (RFE) given by

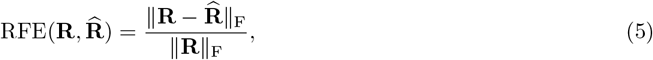

where 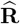 is the approximation of the in-sample LD matrix **R**. This quantity measures the magnitude of the error relative to the magnitude of the full in-sample matrix, providing a scale-invariant metric. However, given the sparse nature of the approximate LD matrices 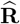, noise from the off-band elements of the in-sample LD will tend to dominate this metric. Therefore, we explored other quantities that focus more narrowly on the error incurred near the diagonal of the LD matrix. For a user-defined bandwidth *b*, we can separate the relative errors within and outside the band as follows:

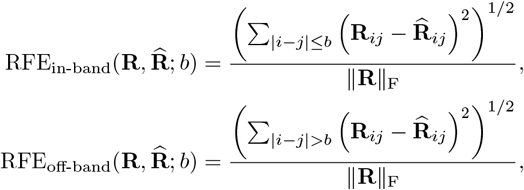

noticing that the square of the each of the two terms sum up to the square of the total RFE defined above, though 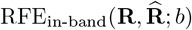 is the main quantity of interest in our analysis. In some cases, it might be useful to restrict the normalizing factor to be localized within the defined band as well:

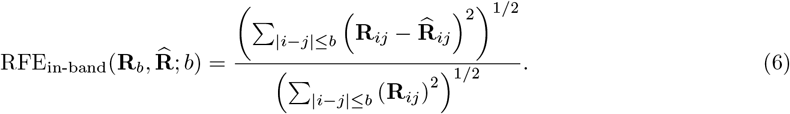

Lastly, we also examined the distribution of individual error terms 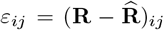 within the band. Also, we used these individual errors to compute the Root Mean Square Error (RMSE), which is equivalent to the unnormalized version of 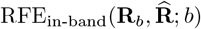. Further analysis of metrics that adjust for sparsity can be found in the Supplementary Section S3.

## Results

To evaluate the performance of our proposed Cholesky-based approach for approximating LD matrices, we compared it against a variety of established methods from the statistical genetics literature. For these evaluations, we used a moderately high-dimensional genotype dataset from The 1000 Genomes Project [37] (*n* = 378 individuals, *m* = 15, 938 SNPs on Chromosome 22) to generate a dense in-sample LD matrix, which served as our ground truth. From this dense matrix, we constructed a number of sparse approximations: a block-diagonal matrix based on LD blocks defined by `LDetect` [21], a shrunk LD matrix based on the shrinkage estimator of Wen and Stephens (2010) [26], and our proposed sparse Cholesky decomposition with fixed bandwidth of 1000 variants and with an adaptive bandwidth of 1.3cM. For reference, we also include a commonly-used banded matrix, with the window size given by the largest block from the previous method. Note that this banded approximation often does not satisfy the positive-definiteness property [16], but we include it here as a baseline for the accuracy metrics since it preserves all terms that are nonzero in either the block-diagonal or Cholesky banded version and is therefore guaranteed to have lower error.

Our analysis first focused on the overall approximation accuracy, quantified by the Relative Frobenius Error (RFE) defined in Equation (5), as shown in Figure 2a. This panel presents the RFE for each approximation method, evaluated across three different bandwidth masks. When compared to the full matrix, all methods exhibit a high total RFE of ∼0.78-0.83. This large error is dominated by the vast number of off-diagonal elements that are set to zero. From this perspective, all sparse methods are comparable as they discard the same long-range information (or noise).

**Figure 2.**
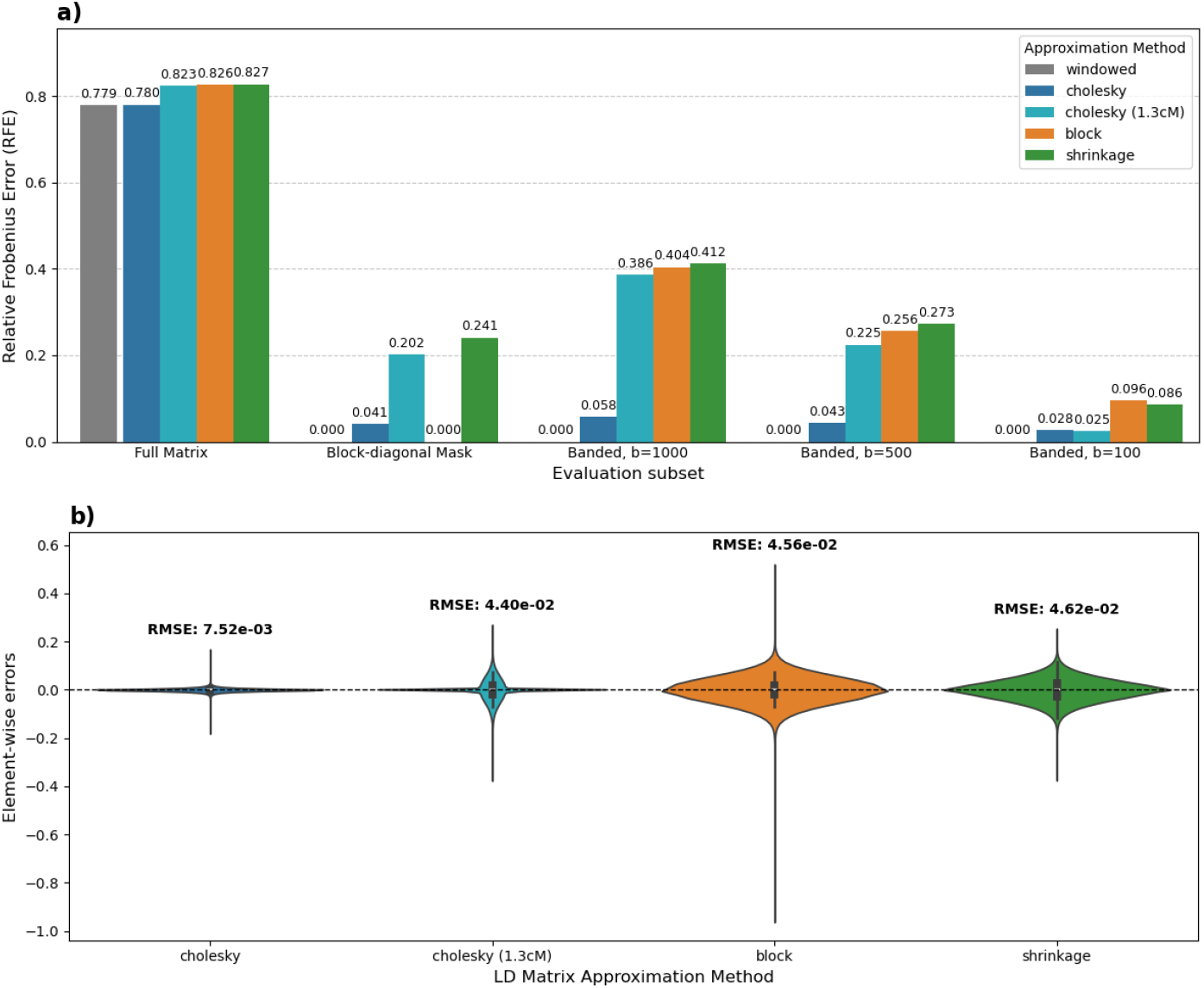
**a)** Performance of approximation methods evaluated across different subsets. The bars show the RFE for each method (indicated by color), calculated within five different masks: the full matrix, the elements within the block-diagonal structure, and banded masks with fixed bandwidth of 1000, 500 and 100 variants. The “windowed” method is set apart to highlight its role as a baseline, as unlike the other methods, it is not guaranteed to be PSD. **b)** Distribution of element-wise errors. Each violin plot shows the probability density of the errors within the window with bandwidth=1000, calculated as estimate - true. The RMSE for each method is annotated above its respective distribution.

The differences between the methods are highlighted by evaluating the error for nearby variants. First, we restrict the calculation to only the elements within the block-diagonal mask, where we use the Frobenius norm of “block” matrix as the denominator when computing the RFE in Equation (5). By their definition, both “windowed” and “block” methods have zero error there. The Cholesky method (based on *b* = 1000) shows a minor relative error of 0.041, while the sparser cM-band Cholesky (1.3cM) method has a higher RFE of 0.202, as its structure is not confined to the LD blocks. The shrinkage approach shows an error of 0.241. Secondly, we evaluate the errors using masks with fixed bandwidth, where we restrict the RFE calculation to only the elements within the band as in Equation (6). Here, the “block” approach incurs a notable RFE of 0.404 ( ∼40%), 0.256 ( ∼26%) and 0.096 ( ∼10%) when using bandwidths given by 1000, 500 and 100 variants, respectively. This indicates that its rigid block structure incorrectly truncates significant correlations that fall within the band but outside the blocks. The shrinkage method shows a similar errors of 0.412, 0.273 and 0.086. The cM-band Cholesky (1.3cM) method shows RFE values of 0.386, 0.225 and 0.025 across the same bands. As shown in the Supplement Section S3, the majority of this error is due to the inherent sparsity of the approximation pattern itself, not a failure to capture LD signals within the defined mask. In contrast, the fixed-band Cholesky method’s RFE of 0.058, 0.043 and 0.028 is substantially lower in both cases, although at the expense of having a denser structure. This demonstrates the key advantage of our approach: it can successfully enforce the PSD property while maintaining the in-band correlation structure.

Next, we investigated the distribution of individual errors within the band to understand the source and magnitude of the approximation errors (Figure 2b). The violin plots reveal the nature of the discrepancies. The block-diagonal method shows a wide error distribution with a heavy tail towards negative values, confirming that its primary source of error is the truncation of strong positive correlations that fall outside its predefined blocks. The shrinkage method has a more symmetric but still wide distribution. In contrast, both Cholesky methods’ error distribution are highly concentrated around zero. This demonstrates that our method not only preserves the overall structure (low RFE) but also accurately maintains the individual correlation values within the band, a crucial feature for downstream fine-mapping and other sensitive analyses.

Finally, we evaluated the computational performance of the method to highlight its scalability to genome-wide data. For these analyses, we used pre-computed block-diagonal LD matrices from Zabad et al. (2025) [16], which are based on genotype data for individuals of European ancestry in the UK Biobank [46]. The largest matrices, corresponding to chromosomes 1 and 2, record pairwise correlations for 116,372 and 119,623 variants, respectively; while the smallest matrices, corresponding to chromosomes 21 and 22, contain data for 20,284 and 20,857 variants, respectively. Detailed information on the size of each chromosome, as well as the amount of non-zero elements within a 1 cM band, can be found in Supplementary Table S2. For the Cholesky method, we used the parallelized implementation over 20 windows, each with its own core, a band-width of 1cM, and a stopping criterion of 3 consecutive iterations with less than 1% relative improvement (see Supplement Section S2 for full hyperparameter details). On the larger chromosomes, the convergence times were approximately 35 minutes, while for the smaller chromosomes the method converged in approximately 6 minutes (Figure 3; Supplementary Table S3). At the same time, the computation of the Cholesky approximation required between 7 and 30GB of memory for most chromosomes (Figure 3; Supplementary Table S3). These results demonstrate empirically that the proposed method can be successfully applied to high-dimensional genomic data, at reasonable computational cost.

**Figure 3.**
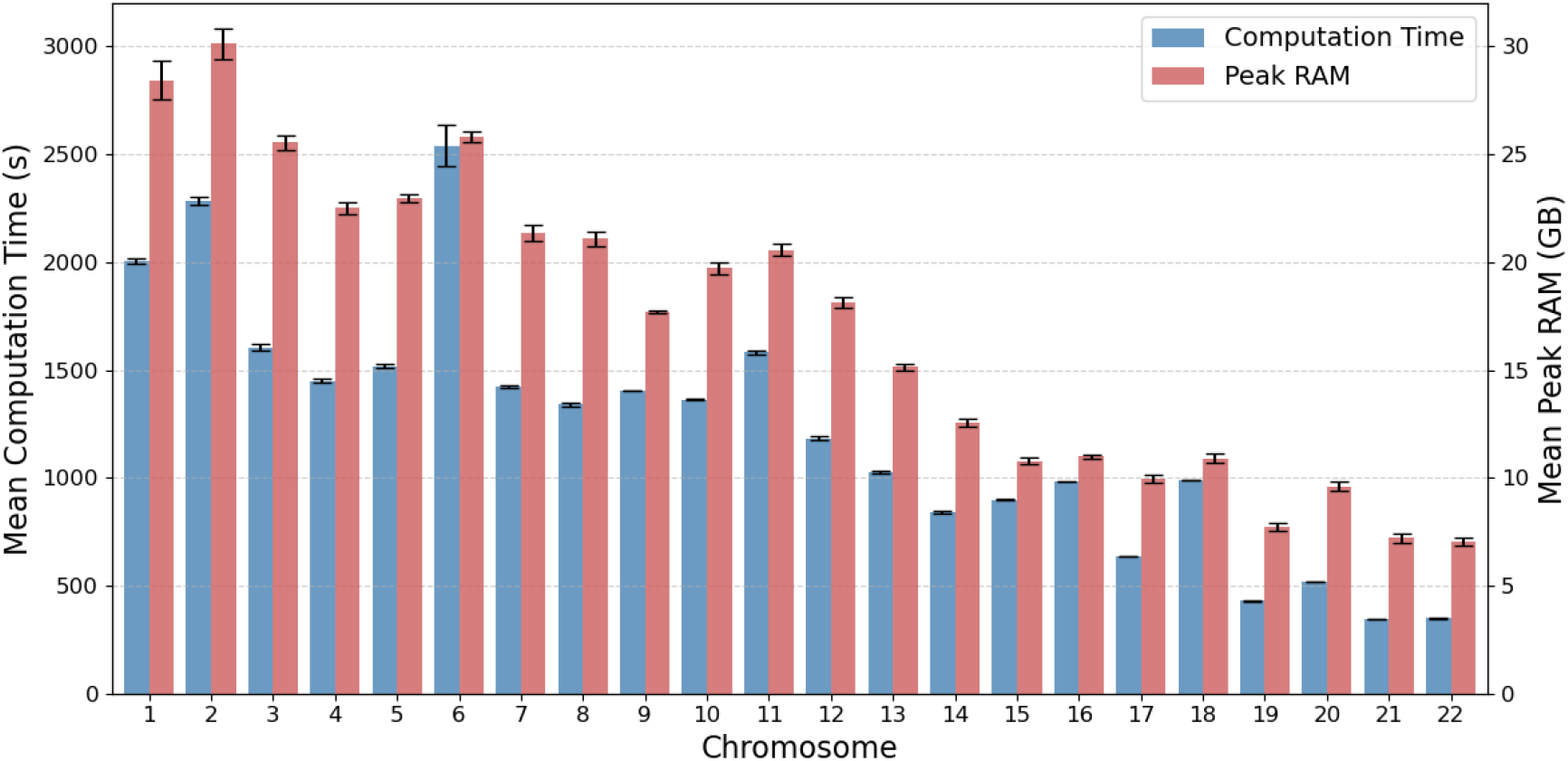
Computational performance of the Cholesky method (1cM band) across the 22 human autosomes, comprising a total of 1.34 million variants. The figure displays the mean computation time in seconds (left blue bars, left axis) and the mean peak RAM (right red bars, right axis) in gigabytes. The values are the averages over five replicates, with error bars representing the standard deviation.

## Discussion

In this work, we introduced a scalable method to address a critical challenge in statistical genetics: generating a sparse LD matrix that is both numerically stable and structurally accurate. The characteristic structure of genomic correlation, where LD is concentrated around the diagonal, allows for efficient, sparse approximations. However, the two most common approaches each have a significant drawback. Banded matrices, which retain all correlations within a fixed distance, often result in an indefinite matrix, compromising the stability of downstream numerical algorithms. Conversely, block-diagonal matrices are guaranteed to be positive semi-definite (PSD), but their rigid block structure can fail to capture significant, high-LD SNP pairs that fall outside the predefined blocks. Our approach is designed to overcome both limitations. We reparameterize the nearest correlation matrix problem via the Cholesky decomposition, which allows us to enforce the PSD property implicitly while retaining the flexible, distance-aware structure of a banded matrix. This avoids the computational bottleneck of repeated eigen-decompositions and yields an approximation that is both numerically stable and structurally faithful to the original data.

To investigate the performance of our method, we compared it against two established techniques for generating PSD approximations—a block-diagonal approach and a shrinkage estimator—using real genotype data from the 1000 Genomes Project. Our analyses, using both the Relative Squared Frobenius Error and the distribution of element-wise errors, showed a marked improvement over these alternatives. The results revealed that while methods like the block-diagonal approximation increases the sparsity and retains the PSD property, they do so at the cost of truncating genuine, high-LD signals that fall outside their rigid block definitions. The Cholesky-based approaches, in contrast, were able to capture these signals with high fidelity.

These analyses demonstrate, in particular, that the PSD property can be recovered for a banded matrix with minimal error without affecting the sparsity. The primary contribution of this work is therefore a method that produces a sparse, computationally efficient, and numerically stable (PSD) approximation of the full LD matrix, enabling more robust downstream analyses such as GWAS fine-mapping and polygenic scoring. Furthermore, we anticipate that the Cholesky factor that our algorithm outputs can be used directly for applications that require sampling from or evaluating multivariate Gaussian likelihoods conditional on the LD matrix [25, 28]. This includes applications such as imputation, simulation, or post-hoc splitting of GWAS summary statistics [25, 47, 48].

Sparse Cholesky decomposition methods are commonly used in the solution of partial differential equations, and many approaches have been proposed to optimize speed and accuracy in that case [49]. However, the tradeoffs are quite different: incomplete Cholesky decomposition for PDEs must be very fast but can tolerate inaccuracies due to further correction mechanisms [49]. By contrast, priority in statistical genomics is accuracy: first, because statistical geneticists typically use the sparse approximations of the LD matrix as the truth, without further correction mechanisms. Second, because LD matrices are usually computed once and re-used for many purposes: some extra work at computation time is warranted. We thus want a method that scales well to large problems, offers accurate representation of the original LD matrix with a reasonable run-time. Here we show that gradient-based minimization of LD residuals performs well.

By developing a gradient descent procedure that leverages the sparse, banded structure of the problem, our implementation gains several practical advantages that enhance its utility. For instance, this design makes the algorithm highly flexible: it can be initialized with existing PSD approximations, such as a block-diagonal matrix, using this information as a “warm start” to accelerate convergence; and allows the use of adaptive bands to account for recombination differences within the genome.

The most significant improvements in accuracy occur within the initial iterations, making the method effective even under time-constrained conditions. Even a few gradient descent steps are guaranteed to return a valid sparse PSD matrix that is no worse than the starting point. Another advantage of the approach presented here is that it is highly flexible and can easily be made to accommodate other desirable properties, such as quantization for more compact storage. With the help of GPU-accelerated automatic differentiation libraries, such as PyTorch/Jax [50], our formulation of the problem can be solved even without analytically tractable gradients, especially once these libraries have more mature interfaces for sparse tensors.

Scalability is critical to large-scale genomic data analysis. The fundamental part of the efficiency of the proposed algorithm lies in the fact that each iteration has a complexity of 𝒪 (*mb*^2^) while allowing parallelization. We can leverage the banded structure to implement a divide-and-conquer strategy to compute the gradient: we can divide the problem into windows such that, to update a window, we only require that region and its adjacent blocks. Consequently, the most intensive computation of the approach, the calculation of the gradient, can be done in parallel without the need for construction or storage of the full matrix. Furthermore, by using the Adam optimizer, we have an approach where the step size is computed locally and efficiently. This allows the method to not be limited by memory while being highly scalable through the use of parallel computation across different cores, which contrast current methods that need the eigen-decomposition [51], the Hessian matrix [32], or computing the inverse of matrices [27]. This was shown by the use of our method on the 22 autosomes, where the convergence time was approximately 33 minutes on the largest chromosome.

Despite its advantages in creating numerically stable and locally accurate LD matrices, our method has limitations: first, like other methods based on banded or block-diagonal structures, our approach discards long-range LD information. While this is a valid tradeoff for many analyses, it may be suboptimal for studying genomic regions with complex, long-range interactions. Recent work has demonstrated the power of combining sparse, local LD matrices with separate, low-rank models that specifically capture long-range LD [27]. Second, the accuracy of any in-sample LD matrix, including our approximation, is based on the quality of the input genotype data. Our method operates on the provided LD matrix, but does not itself correct for biases that may be present in it before our algorithm is applied. Major sources of such bias are population structure [52, 53] and small sample size [6, 17, 54]. Therefore, for our method to yield the most accurate representation of population LD, it is recommended that the input genotype data be pre-processed to account for these factors.

## Code and data availability

The Python code implementing the “cholesky” algorithm, as well as the code used to generate the figures and tables in this study, are publicly available in our `GitHub` repository at https://github.com/uliBercovich/SPLD.

## Acknowledgments

We thank Anton Rask Lundborg for helpful discussions regarding gradient descent on Riemannian manifolds.

## Conflicts of interests

The authors have no conflicts of interest to declare.

## Funding

This research was supported by the Canadian Institute for Health Research (CIHR) project grant 437576, NSERC grant RGPIN-2023-04882; the Canada Research Chair program to S.G.; the Canada Foundation for Innovation; and the Independent Research Fund Denmark (grant number: DFF-8021-00360B) to U.B.

## Notes

### Competing Interest Statement

The authors have declared no competing interest.

## References

1. Pritchard, J. K. & Przeworski, M. Linkage Disequilibrium in Humans: Models and Data. The American Journal of Human Genetics 69, 1–14 (2001).

2. Slatkin, M. Linkage disequilibrium —understanding the evolutionary past and mapping the medical future. Nature Reviews Genetics 9, 477–485 (2008).

3. Sved, J. A. & Hill, W. G. One Hundred Years of Linkage Disequilibrium. Genetics 209, 629–636 (2018).

4. Hill, W. G. & Robertson, A. Linkage disequilibrium in finite populations. Theoretical and Applied Genetics 38, 226–231 (1968).

5. Tishkoff, S. A. et al. Global Patterns of Linkage Disequilibrium at the CD4 Locus and Modern Human Origins. Science 271, 1380–1387 (1996).

6. Ragsdale, A. P. & Gravel, S. Unbiased Estimation of Linkage Disequilibrium from Unphased Data. Molecular Biology and Evolution 37, 923–932 (2019).

7. Ragsdale, A. P. et al. A weakly structured stem for human origins in Africa. Nature 617, 755–763 (2023).

8. Devlin, B. & Risch, N. A Comparison of Linkage Disequilibrium Measures for Fine-Scale Mapping. Genomics 29, 311–322 (1995).

9. Risch, N. & Merikangas, K. The Future of Genetic Studies of Complex Human Diseases. Science 273, 1516–1517 (1996).

10. Altshuler, D., Daly, M. J. & Lander, E. S. Genetic Mapping in Human Disease. Science 322, 881–888 (2008).

11. Chen, S. et al. A genomic mutational constraint map using variation in 76,156 human genomes. Nature 625, 92–100 (2024).

12. Li, S., Carss, K. J., Halldorsson, B. V. & Cortes, A. Whole-genome sequencing of half-a-million UK Biobank participants. medRxiv (2023).

13. Schaid, D. J., Chen, W. & Larson, N. B. From genome-wide associations to candidate causal variants by statistical fine-mapping. Nature Reviews Genetics 19, 491–504 (2018).

14. Wang, Y. et al. Theoretical and empirical quantification of the accuracy of polygenic scores in ancestry divergent populations. Nature Communications 11, 3865 (2020).

15. Zheng, Z. et al. Leveraging functional genomic annotations and genome coverage to improve polygenic prediction of complex traits within and between ancestries. Nature Genetics 56, 767–777 (2024).

16. Zabad, S., Haryan, C. A., Gravel, S., Misra, S. & Li, Y. Toward whole-genome inference of polygenic scores with fast and memory-efficient algorithms. The American Journal of Human Genetics 112, 1528–1546 (2025).

17. Bulik-Sullivan, B. et al. LD score regression distinguishes confounding from polygenicity in genome-wide association studies. Nature Genetics 47 (3 2015).

18. Mak, T. S. H., Porsch, R. M., Choi, S. W., Zhou, X. & Sham, P. C. Polygenic scores via penalized regression on summary statistics. Genetic Epidemiology 41 (6 2017).

19. Lloyd-Jones, L. R. et al. Improved polygenic prediction by Bayesian multiple regression on summary statistics. Nature Communications 10 (1 2019).

20. Pasaniuc, B. & Price, A. L. Dissecting the genetics of complex traits using summary association statistics. Nature Reviews Genetics 18, 117–127 (2017).

21. Berisa, T. & Pickrell, J. K. Approximately independent linkage disequilibrium blocks in human populations. Bioinformatics 32, 283–285 (2015).

22. Privé, F., Arbel, J. & Vilhjálmsson, B. J. LDpred2: Better, faster, stronger. Bioinformatics 36 (22–23 2020).

23. Privé, F. Optimal linkage disequilibrium splitting. Bioinformatics 38, 255–256 (2021).

24. Zabad, S., Gravel, S. & Li, Y. Fast and accurate Bayesian polygenic risk modeling with variational inference. The American Journal of Human Genetics 110, 741–761 (2023).

25. Salehi Nowbandegani, P. et al. Extremely sparse models of linkage disequilibrium in ancestrally diverse association studies. Nature Genetics 55, 1494–1502 (2023).

26. Wen, X. & Stephens, M. Using linear predictors to impute allele frequencies from summary or pooled genotype data. Ann Appl Stat 4, 1158–1182 (2010).

27. Li, H., Mazumder, R. & Lin, X. Accurate and efficient estimation of local heritability using summary statistics and the linkage disequilibrium matrix. Nature Communications 14, 7954 (2023).

28. Benner, C. et al. FINEMAP: efficient variable selection using summary data from genome-wide association studies. Bioinformatics 32, 1493–1501 (2016).

29. Benner, C. et al. Prospects of Fine-Mapping Trait-Associated Genomic Regions by Using Summary Statistics from Genome-wide Association Studies. The American Journal of Human Genetics 101, 539–551 (2017).

30. Higham, N. J. Computing the nearest correlation matrix—a problem from finance. IMA Journal of Numerical Analysis 22, 329–343 (2002).

31. Knol, D. L. & ten Berge, J. M. F. Least-Squares Approximation of an Improper Correlation Matrix by a Proper One. Psychometrika 54, 53–61 (1989).

32. Lurie, P. M. & Goldberg, M. S. An Approximate Method for Sampling Correlated Random Variables from Partially-Specified Distributions. Management Science 44, 203–218 (1998).

33. Lucas, C. Computing Nearest Covariance and Correlation Matrices M.Sc. Thesis (University of Manchester, 2001).

34. Qi, H. & Sun, D. A Quadratically Convergent Newton Method for Computing the Nearest Correlation Matrix. SIAM Journal on Matrix Analysis and Applications 28, 360–385 (2006).

35. Journée, M., Bach, F.Absil, P.-A. & Sepulchre, R. Low-Rank Optimization on the Cone of Positive Semidefinite Matrices. SIAM Journal on Optimization 20, 2327–2351 (2010).

36. Benoit & Cholesky. Note Sur Une Méthode de Résolution des équations Normales Provenant de L’Application de la MéThode des Moindres Carrés a un Système D’équations Linéaires en Nombre Inférieur a Celui des Inconnues. — Application de la Méthode a la Résolution D’un Système Defini D’éQuations LinéAires. Bulletin géodésique 2, 67–77 (1924).

37. Consortium, T. 1. G. P. A global reference for human genetic variation. Nature 526, 68–74 (2015).

38. Absil, P., Mahony, R. & Sepulchre, R. Optimization Algorithms on Matrix Manifolds (Princeton University Press, 2008).

39. Kim, S. A., Cho, C.-S., Kim, S.-R., Bull, S. B. & Yoo, Y. J. A new haplotype block detection method for dense genome sequencing data based on interval graph modeling of clusters of highly correlated SNPs. Bioinformatics 34, 388–397 (2017).

40. George, A. & Liu, J. W. H. Computer Solution of Large Sparse Positive Definite Systems (Prentice Hall, 1981).

41. Armijo, L. Minimization of functions having Lipschitz continuous first partial derivatives. Pacific Journal of Mathematics 16, 1–3 (1966).

42. Goldstein, A. A. On Steepest Descent. Journal of the Society for Industrial and Applied Mathematics Series A Control 3, 147–151 (1965).

43. Kingma, D. P. & Ba, J. Adam: A Method for Stochastic Optimization 2017.

44. Harris, C. R. et al. Array programming with NumPy. Nature 585, 357–362 (2020).

45. Virtanen, P. et al. SciPy 1.0: Fundamental Algorithms for Scientific Computing in Python. Nature Methods 17, 261–272 (2020).

46. Bycroft, C. et al. The UK Biobank resource with deep phenotyping and genomic data. Nature 562 (7726 2018).

47. Pasaniuc, B. et al. Fast and accurate imputation of summary statistics enhances evidence of functional enrichment. Bioinformatics 30, 2906–2914 (July 2014).

48. Zhao, Z. et al. PUMAS: fine-tuning polygenic risk scores with GWAS summary statistics. Genome Biology 22, 257 (Sept. 6, 2021).

49. Benzi, M. Preconditioning Techniques for Large Linear Systems: A Survey. Journal of Computational Physics 182, 418–477 (2002).

50. Paszke, A. et al. PyTorch: An Imperative Style, High-Performance Deep Learning Library in Advances in Neural Information Processing Systems 32 8024–8035 (Curran Associates, Inc., 2019).

51. Borsdorf, R. & Higham, N. J. A preconditioned Newton algorithm for the nearest correlation matrix. IMA Journal of Numerical Analysis 30, 94–107 (2009).

52. Mangin, B. et al. Novel measures of linkage disequilibrium that correct the bias due to population structure and relatedness. Heredity 108, 285–291 (2012).

53. Bercovich, U., Rasmussen, M. S., Li, Z., Wiuf, C. & Albrechtsen, A. Measuring linkage disequilibrium and improvement of pruning and clumping in structured populations. Genetics 229, iyaf009 (Feb. 2025).

54. Waples, R. S. A bias correction for estimates of effective population size based on linkage disequilibrium at unlinked gene loci. Conservation Genetics 7, 167–184 (2006).

