## Supplementary material for "LD Matrix Approximations for Scalable Analysis of High-dimensional Genetic Data"

#### S1 Supplementary Theory

**Definition S1.** *For a matrix* $\mathbf{M}$*, we say that* $\mathbf{M}$ *has bandwidth* $b$ *if* $\mathbf{M}_{ij} = 0$ *for every pair* $i$*,* $j$ *such that* $|i - j| > b$*.*

The following result connects the bandwidth of $\mathbf{X}$ with the bandwidth of $\mathbf{L}$, where $\mathbf{L}$ is such that $\mathbf{X} = \mathbf{L}\mathbf{L}^{\mathrm{T}}$. A proof can be found in Theorem 4.3.1 in [1]. A general case using more flexible patterns of sparsity can be found in Lemma 4.2.1 in [2].

**Lemma S1.**
1. *Let* $\mathbf{X}$ *be a positive semi-definite (PSD) matrix with bandwidth* $b$ *such that* $\mathbf{X} = \mathbf{L}\mathbf{L}^{\mathrm{T}}$*. Then,* $\mathbf{L}$ *has bandwidth* $b$*.*
2. *Let* $\mathbf{L}$ *be a lower triangular matrix with bandwidth* $b$*. Then,* $\mathbf{X} = \mathbf{L}\mathbf{L}^{\mathrm{T}}$ *has bandwidth* $b$*.*

We can now proceed at proving how the feasible set of Problem (4) provides a parametrization of the feasible set of Problem (3).

**Lemma S2.**
1. *Let* $\mathbf{X}^*$ *be feasible solution of Problem* (3)*. Then, there is a feasible solution* $\mathbf{L}^*$ *of Problem* (4) *such that* $g(\mathbf{L}^*) = \mathbf{X}^*$*.*
2. *Let* $\mathbf{L}^*$ *be feasible solution of Problem* (4)*. Then,* $g(\mathbf{L}^*) = \mathbf{X}^*$ *is a feasible solution of Problem* (3)

*Proof.*
1. Let $\mathbf{L}^*$ be the generalized Cholesky decomposition of $\mathbf{X}^*$, which exist as $\mathbf{X}^*$ is PSD. By Lemma S1, $\mathbf{L}^*$ has the same bandwidth as $\mathbf{X}^*$. As each diagonal element of $\mathbf{X}^*$ is 1, we have that $\mathbf{L}^*(\mathbf{L}^*)^{\mathrm{T}}$ is equal to 1 on each element of the diagonal, which is equivalent to saying that $\|\mathbf{L}_{i\cdot}\|_2^2 = 1$ for every $i = 1, \ldots, m$. Finally, by definition of the Cholesky decomposition, $\mathbf{L}^*$ is lower triangular, satisfying every requirement of Problem (4)

2. By definition, $g(\mathbf{L}^*) \succeq 0$. By Lemma S1, $g(\mathbf{L}^*)$ has the same bandwidth as $\mathbf{L}^*$. Furthermore, the diagonal constrains are satisfied by the fact that $(g(\mathbf{L}^*))_{ii} = \|\mathbf{L}^*_{i\cdot}\|_2^2 = 1$ for every $i = 1, \ldots, m$. Hence, $g(\mathbf{L}^*)$ is a feasible solution for Problem (3).

□

#### S2 Comparison of strategies to estimate the step size

A comparison between the two strategies to select the step size on the Cholesky-based approach was also performed. Both implementations were done on `Python` using `numpy` [3] and `scipy.sparse` for sparse matrix computations, and `joblib` [4] for parallelization. For the backtracking approach, a warm start based on doubling the previous step size was used, with standard parameters given by the initial step size $\alpha = 1$, the contraction parameter $\beta = 0.5$, and the Armijo parameter $c = 10^{-4}$. For the Adam approach, we explored different learning rates while keeping recommended parameters fixed ($\beta_1 = 0.9$, $\beta_2 = 0.999$, $\varepsilon = 10^{-8}$), but kept the recommended learning rate of 0.001 [5]. For parallelization, we used 20 windows processed on 20 CPU cores. The stopping criteria was a relative improvement of less than 0.1% over ten consecutive iterations. Figure S1 illustrates the superior performance of the parallelized Adam optimizer over the standard backtracking line search. Across all tested matrix sizes, the Adam approach was consistently faster and achieved a lower final approximation error. For the largest matrix ($m$=15,937), the Adam optimizer provided a speedup of one order of magnitude (40 minutes vs. 6 hours 34 minutes) while also reaching a better solution (Squared Frobenius Error of $2.28 \times 10^3$ vs. $2.96 \times 10^3$). This demonstrates that the parallel Adam approach is both more efficient and more effective for this high-dimensional optimization problem.

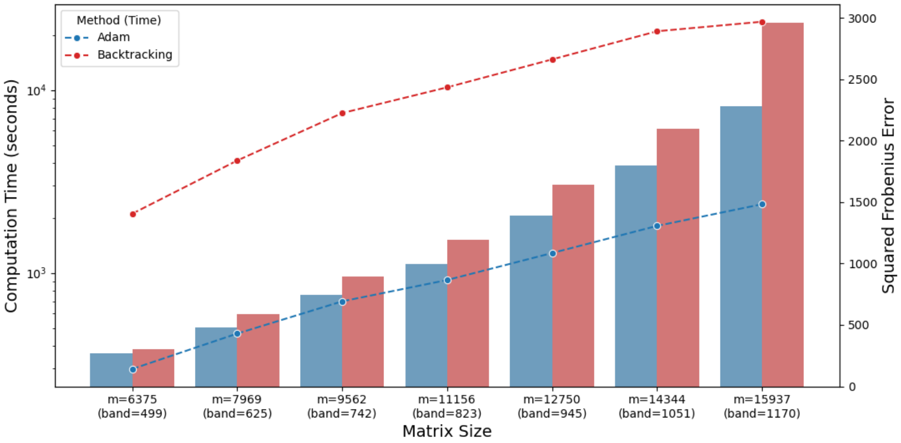


Figure S1: Comparison of computational performance and accuracy for the Backtracking and Adam methods across increasing problem sizes. Computation time (in seconds) is plotted on the primary y-axis (left, logarithmic scale) and represented by dashed lines with markers. The corresponding Squared Frobenius Error ($\|\mathbf{R} - \hat{\mathbf{R}}\|_{\mathrm{F}}^2$) is plotted on the secondary y-axis (right) and represented by bars. Each category on the x-axis corresponds to a specific matrix dimension (m) and bandwidth (b).

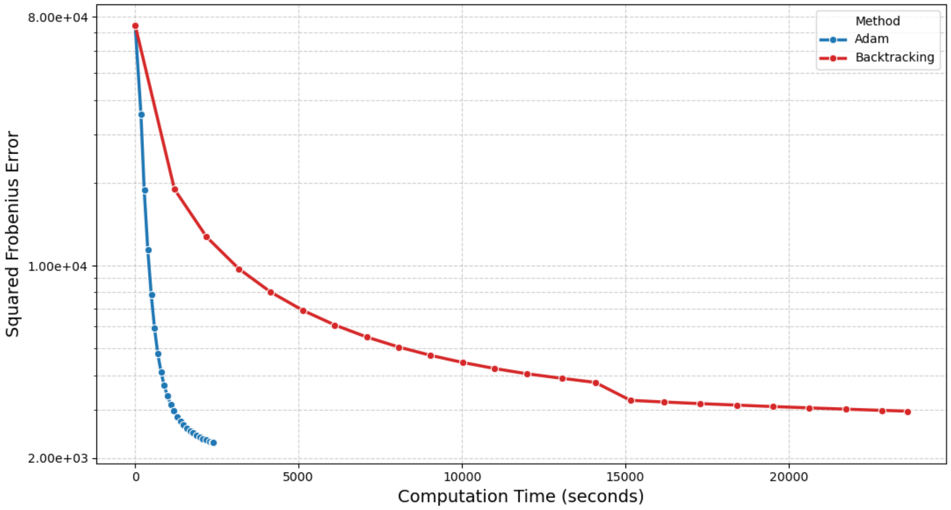


Figure S2: Comparison of the backtracking and Adam methods. The plot shows the decrease in Squared Frobenius Error ($\|\mathbf{R} - \hat{\mathbf{R}}\|_{\mathrm{F}}^2$) as a function of wall-clock computation time in seconds. Each data point represents a measurement during the optimization process every ten iterations. The error axis is presented on a logarithmic scale.

#### S3 Sparsity aware Frobenius metric

We are interested in estimating the loss of LD information caused by the different sparsity assumption of the different methods. Let $\varepsilon_{ij}$ be the sample correlation between markers such that $|i-j|$ is large. Hence, by taking distant SNPs, we can assume that the LD on a population level is 0. Furthermore, we assume that this error terms $\varepsilon_{ij}$ have a normal distribution with parameters $(0, \sigma^2)$ (Figure S4). Then, by using the Law of Large Numbers, we define the expected off-band relative Frobenius error as

$$\widetilde{\mathrm{RFE}}_{\text{off-band}}(\mathbf{R}; b) := \frac{\sqrt{(m^2 - N_b)}\sigma}{\|\mathbf{R}\|_{\mathrm{F}}} \approx \frac{\sqrt{\sum_{|i-j|>b} \varepsilon_{ij}}}{\|\mathbf{R}\|_{\mathrm{F}}},$$

where $N_b$ is the amount of elements within the band. This $\widetilde{\mathrm{RFE}}_{\text{off-band}}(\mathbf{R})$ can be understood as the error that we cannot diminish under the assumption that the elements outside the band are only noise without LD signals. This can be further extended to any mask $\mathcal{M}$ of $m \times m$ to define

$$\widetilde{\mathrm{RFE}}_{\mathcal{M}}(\mathbf{R}) = \frac{\sqrt{\sum_{(i,j)\notin\mathcal{M}} \sigma^2}}{\|\mathbf{R}\|_{\mathrm{F}}} = \frac{\sqrt{(m^2 - |\mathcal{M}|)}\sigma}{\|\mathbf{R}\|_{\mathrm{F}}},$$

where $|\mathcal{M}|$ is the amount of non-zero elements of the mask. Furthermore, we can focalize on the in-band section by using the banded $\mathbf{R}_b$ to compute

$$\widetilde{\mathrm{RFE}}_{\text{in-band},\mathcal{M}}(\mathbf{R}_b; b) = \frac{\sqrt{(N_b - |\mathcal{M}|)}\sigma}{\|\mathbf{R}_b\|_{\mathrm{F}}},$$

for a mask $\mathcal{M}$ within the band.

Then, given a banded estimate $\widehat{\mathbf{R}}$ within the mask $\mathcal{M}_{\text{band}}$, a larger $\mathrm{RFE}(\widehat{\mathbf{R}})$ than the expected $\widetilde{\mathrm{RFE}}_{\mathcal{M}_{\text{band}}}(\mathbf{R})$ would mean that there are errors that are purely caused by the estimate. In this way, we can discriminate between errors from the estimation and errors given by the inevitable lack of coverage of noise terms, which is due to the sparse nature of the method. Then, the value of interest will be the difference $\mathrm{RFE}(\widehat{\mathbf{R}}) - \widetilde{\mathrm{RFE}}_{\mathcal{M}_{\text{band}}}(\mathbf{R})$, i.e., the estimate errors that are not explained by the background noise or, to frame it differently, the high-LD signals that are lost by the sparsification.

This metric can be extended to the block-diagonal sparsification: let $\mathcal{M}_{\text{block}}$ be the block-diagonal mask applied by this method. Then,

$$\widetilde{\mathrm{RFE}}_{\mathcal{M}_{\text{block}}}(\mathbf{R}) = \frac{\sqrt{(m^2 - |\mathcal{M}_{\text{block}}|)}\sigma}{\|\mathbf{R}\|_{\mathrm{F}}}.$$

This can help us to make a sparse-aware comparison between different methods that work with a predefined mask. In particular, by using this metric, we isolate the error from the dominant off-band noise, an effect that is further enhanced when utilizing $\widetilde{\mathrm{RFE}}_{\text{in-band},\mathcal{M}}(\mathbf{R}_b; b)$.

In Figure S3, we compare the sparser "cholesky" using an adaptive band of 1.3 centiMorgan (cM) and the "block" methods. As "shrinkage" is not based on a predefined sparse structure, it is excluded from this analysis. To estimate $\sigma$, we estimated the variance of the off-band elements as in Figure S4 (bottom right panel). Panel a) reproduces the total RFE from Figure 2 with the addition of the expected RFE (dashed lines). For the adaptive-band Cholesky and the block-diagonal approach, it estimates the error that we should have on the case where every element not covered by a block is pure noise. Panel b) computes the difference of RFE - expected RFE, i.e. the estimation error that is not an effect of the sparsification.

On the Cholesky approach, we see a decrease of the error terms showing that the numerical cost of enforcing the PSD property is primarily absorbed by correlations further away from the diagonal. Moreover, the adjusted errors on panel b) are heavily diminished, which is a result of the correctly ignored noise terms dominating the plain RFE metric of panel a). In contrast, we observe the opposite trend on the block-diagonal method, where the narrower the band is, the higher is the difference between the RFE and the expected RFE. We can see that on a larger band, the noise terms dominates the lost LD signals, making both the RFE and the expected RFE similar, as with the sparser Cholesky approach. However, when we focus in correlations from close markers by using a smaller bandwidth, we have the opposite effect: within a narrower band that is dense with true LD signals, the structural error from incorrectly truncated correlation becomes the dominant component. This confirms that the block-diagonal structure is unable to capture high LD pairs between variants that are not covered by the predefined blocks. Therefore, the closer the variants are, the better the Cholesky method is at keeping the LD signals, an effect not sustained by the block-diagonal approach.

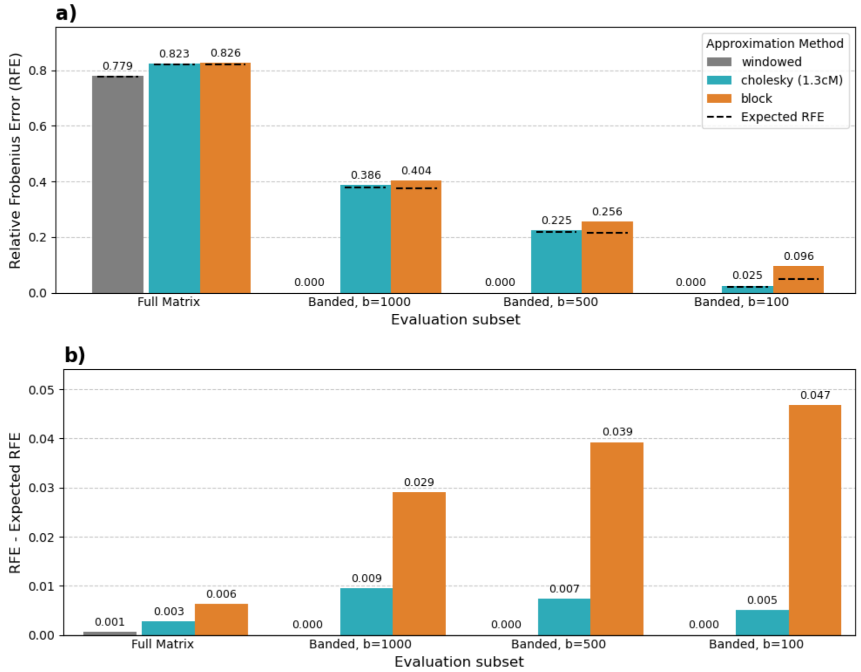


Figure S3: **a)** Performance of approximation methods evaluated across different subsets. The bars show the RFE for each method (indicated by color), calculated within four different masks: the full matrix, and banded masks with fixed bandwidth of 1000, 500 and 100 variants. The dashed line indicates the expected $\widetilde{\mathrm{RFE}}_{\text{in-band},\mathcal{M}}(\mathbf{R}_b; b)$ for each method and band. **b)** Adjusted RFE, calculated as the difference between the total RFE and the Expected RFE. This metric isolates the error that is attributable to the approximation algorithm itself.

#### Supplementary algorithms

Pseudo code of both search methods for selecting $\alpha$ on the Cholesky based approximation are presented. Algorithm S1 presents the backtracking approach, Algorithm S2 presents the one using Adam, with Algorithm S3 an auxiliary function used in Adam.

**Algorithm S1** Nearest Banded Correlation Matrix with Backtracking Line Search

**Input:** Symmetric banded matrix $\mathbf{R}_b$, bandwidth $b$, starting matrix $\mathbf{L}_0$, stopping criteria $k_{max}$ and $\varepsilon$, backtracking parameters $\alpha_0, c, \rho, \varepsilon_\alpha$.

**Output:** Banded and lower triangular matrix $\mathbf{L}$.

$k \leftarrow 0$
$\mathbf{L}_k \leftarrow \Pi_{\mathcal{F}_{Chol}}(\mathbf{L}_0)$ ▷ Project onto the manifold
$f_k \leftarrow \|\mathbf{R}_b - \mathbf{L}_k\mathbf{L}_k^{\mathrm{T}}\|_{\mathrm{F}}^2$
$\alpha \leftarrow \alpha_{init}$ ▷ Initialize step size
**while** $k < k_{\max}$ **do**
  $\nabla f_k \leftarrow \Pi_{\mathcal{S}^1}(\Pi_{\mathcal{L}_b}(4(\mathbf{L}_k\mathbf{L}_k^T - \mathbf{R}_b)\mathbf{L}_k)(\mathbf{L}_k)$ ▷ Compute the Riemannian gradient
  is_acceptable ← FALSE
  **while not** is_acceptable **do**
    $\mathbf{L}_{k+1} \leftarrow \Pi_{\mathcal{F}_{Chol}}(\mathbf{L}_k - \alpha \cdot \nabla f_k)$ ▷ Propose a step
    $f_{k+1} \leftarrow \|\mathbf{R}_b - \mathbf{L}_{k+1}\mathbf{L}_{k+1}^{\mathrm{T}}\|_{\mathrm{F}}^2$
    **if** $f_{k+1} \leq f_k - c \cdot \alpha \cdot \|\nabla f_k\|_{\mathrm{F}}^2$ **then** ▷ Armijo condition
      is_acceptable ← TRUE
    **else**
      $\alpha \leftarrow \rho \cdot \alpha$ ▷ Decrease step size
      **if** $\alpha < \varepsilon_\alpha$ **then** ▷ Step size too small
        **break**
      **end if**
    **end if**
  **end while**
  **if** $(f_k - f_{k+1})/f_k < \varepsilon$ **then** ▷ Check for convergence
    **break**
  **end if**
  $\alpha \leftarrow \alpha/\rho$ ▷ Warm-start next iteration with one larger step
  $k \leftarrow k + 1$
**end while**
$\mathbf{L} \leftarrow \mathbf{L}_k$ ▷ Return the final matrix

**Algorithm S2** Window-based Adam for Nearest Banded Correlation Matrix

**Input:** Symmetric and banded matrix $\mathbf{R}_b$, bandwidth $b$, starting matrix $\mathbf{L}_0$, stopping criteria $k_{max}$ and $\varepsilon$, Adam parameters $\eta, \beta_1, \beta_2, \epsilon$, parallelization parameter $n_{windows}$.

**Output:** Banded and lower triangular matrix $\mathbf{L}$.

$k \leftarrow 0$

$\mathbf{L}_k \leftarrow \Pi_{\mathcal{F}_{Chol}}(\mathbf{L}_0)$ ▷ Project onto the manifold

$f_k \leftarrow \|\mathbf{R}_b - \mathbf{L}_k \mathbf{L}_k^{\mathrm{T}}\|_{\mathrm{F}}^2$

$m_k, v_k \leftarrow \mathbf{0}, \mathbf{0}$ ▷ Initialize 1st and 2nd moment vectors as zero matrices

$W_1, \ldots, W_{n_{windows}} \leftarrow \text{PartitionRows}(m, n_{windows})$ ▷ Divide row indices into windows

**while** $k < k_{\max}$ **do**

  **for** $j \in \{1, \ldots, n_{windows}\}$ **do in parallel**

    $(\mathbf{L}_{k+1}^{(j)}, m_{k+1}^{(j)}, v_{k+1}^{(j)}) \leftarrow \text{WindowUpdate}(W_j, \mathbf{R}_b, b, \mathbf{L}_k, m_k, v_k, k, \eta, \beta_1, \beta_2, \epsilon)$

  **end for**

  $\mathbf{L}_{k+1} \leftarrow \text{AssembleRows}(\mathbf{L}_{k+1}^{(1)}, \ldots, \mathbf{L}_{k+1}^{(n_{windows})})$

  $m_{k+1} \leftarrow \text{AssembleRows}(m_{k+1}^{(1)}, \ldots, m_{k+1}^{(n_{windows})})$

  $v_{k+1} \leftarrow \text{AssembleRows}(v_{k+1}^{(1)}, \ldots, v_{k+1}^{(n_{windows})})$

  $f_{k+1} \leftarrow \|\mathbf{R}_b - \mathbf{L}_{k+1} \mathbf{L}_{k+1}^{\mathrm{T}}\|_{\mathrm{F}}^2$

  **if** $(f_k - f_{k+1})/f_k < \varepsilon$ **then** ▷ Check for convergence

    **break**

  **end if**

  $k \leftarrow k + 1$

**end while**

$\mathbf{L} \leftarrow \mathbf{L}_k$ ▷ Return the final matrix

**Algorithm S3** Worker Function: WindowUpdate

**Input:** Window $W_j$, symmetric and banded matrix $\mathbf{R}_b$, bandwidth $b$, current matrices $\mathbf{L}_k, m_k, v_k$, iteration $k$, Adam parameters $\eta, \beta_1, \beta_2, \epsilon$.

**Output:** Updated window matrices $\mathbf{L}_{k+1}^{\text{win}}, m_{k+1}^{\text{win}}, v_{k+1}^{\text{win}}$.

**function** $\textsc{WindowUpdate}(W_j, \mathbf{R}_b, b, \mathbf{L}_k, \dots)$

  $\mathbf{L}_k^{\text{win}} \leftarrow \mathbf{L}_k[W_j, :]$ ▷ Get rows corresponding to the current window

  $m_k^{\text{win}} \leftarrow m_k[W_j, :]$

  $v_k^{\text{win}} \leftarrow v_k[W_j, :]$

  $\nabla f_k^{\text{win}} \leftarrow \Pi_{\mathcal{S}^1}(\Pi_{\mathcal{L}_b}(4(\mathbf{L}_k^{\text{win}}\mathbf{L}_k^T - \mathbf{R}_b)\mathbf{L}_k)(\mathbf{L}_k^{\text{win}})$ ▷ Compute the Riemannian gradient

  $m_{k+1}^{\text{win}} \leftarrow \beta_1 m_k^{\text{win}} + (1-\beta_1)\nabla f_k^{\text{win}}$

  $v_{k+1}^{\text{win}} \leftarrow \beta_2 v_k^{\text{win}} + (1-\beta_2)(\nabla f_k^{\text{win}} \odot \nabla f_k^{\text{win}})$ ▷ $\odot$: Hadamard product

  $\hat{m}^{\text{win}} \leftarrow m_{k+1}^{\text{win}}/(1-\beta_1^{k+1})$

  $\hat{v}^{\text{win}} \leftarrow v_{k+1}^{\text{win}}/(1-\beta_2^{k+1})$

  $\Delta_{\mathbf{L}}^{\text{win}} \leftarrow \eta \cdot \hat{m}^{\text{win}} \oslash (\sqrt{\hat{v}^{\text{win}}} + \epsilon)$ ▷ Compute Adam update step. $\oslash$: element-wise division

  $\mathbf{L}_{k+1}^{\text{win}} \leftarrow \Pi_{\mathcal{F}_{Chol}}(\mathbf{L}_k^{\text{win}} - \Delta_{\mathbf{L}}^{\text{win}})$ ▷ Projection step

  **return** $(\mathbf{L}_{k+1}^{\text{win}}, m_{k+1}^{\text{win}}, v_{k+1}^{\text{win}})$

**end function**

### Supplementary figures

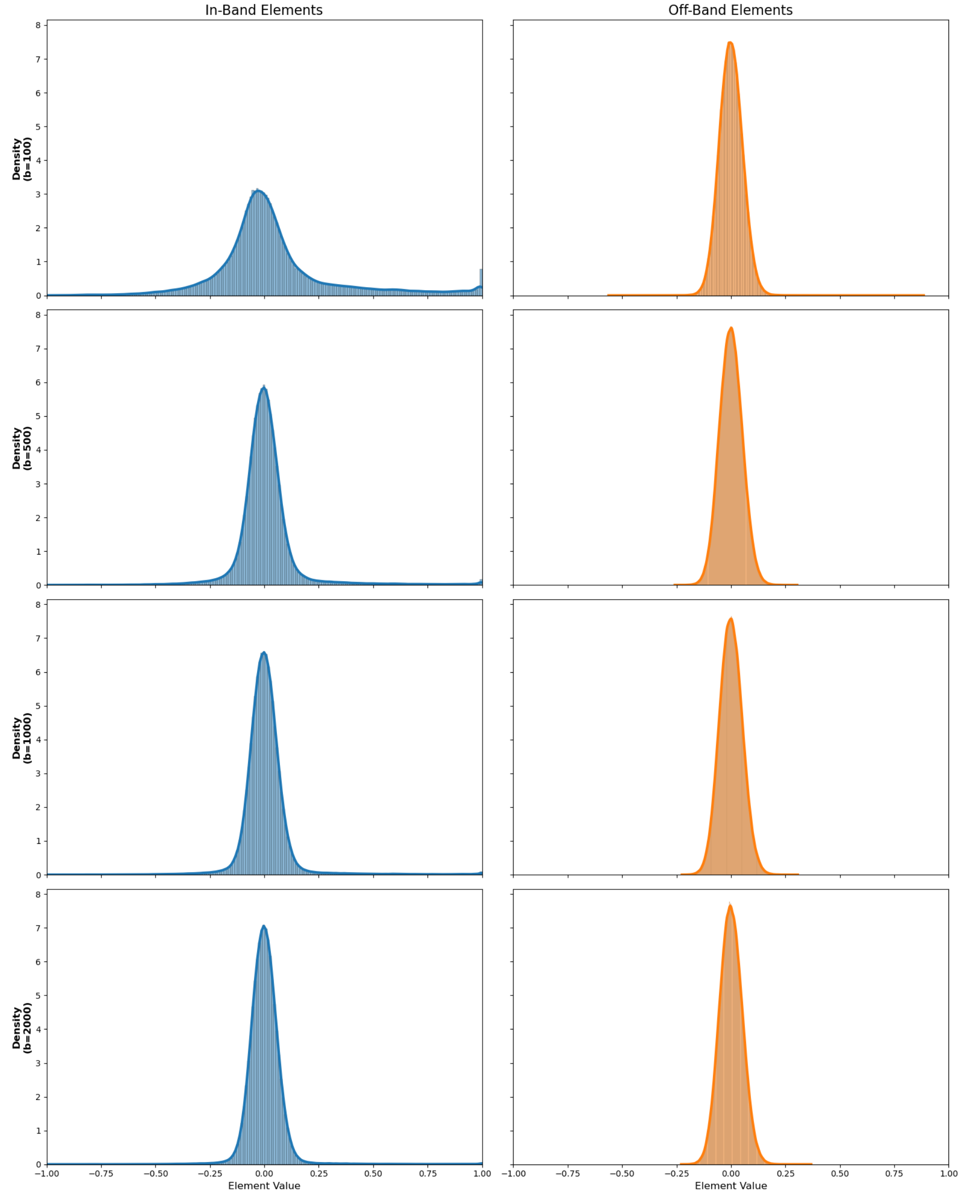


Figure S4: Approximate distribution of in-band and off-band LD matrix elements for varying bandwidths. The left column (blue) shows the distribution of elements inside the band while the right column (orange) shows the distribution of elements outside the band. Each row corresponds to a different bandwidth, increasing from top to bottom (b = 100, 500, 1000, 2000).

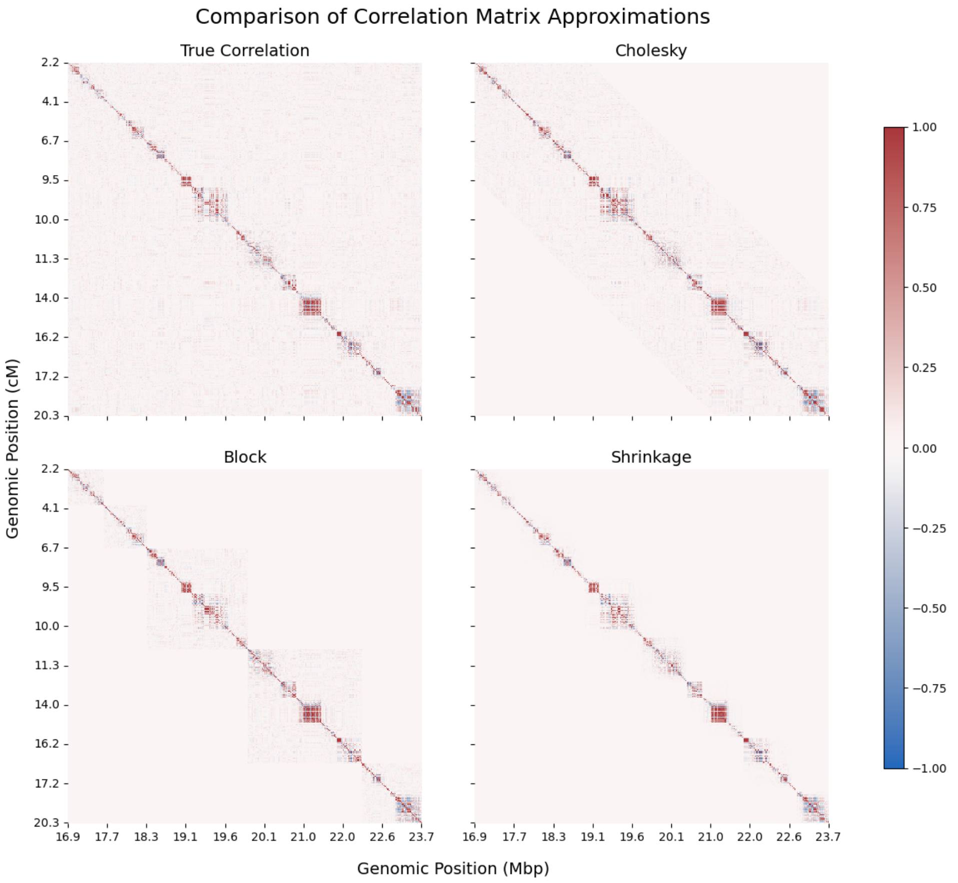


Figure S5: Heatmap visualizations of the ground truth correlation matrix and three approximations for a representative genomic region on Chromosome 22. Top left panel shows the $m \times m$ Pearson correlation matrix $\mathbf{R}$, serving as the ground truth. Top right panel shows the output of our proposed Cholesky-based algorithm, which finds the nearest positive semi-definite (PSD) matrix with a banded structure. Bottom left panel shows the block-diagonal approximation, where correlations between pre-defined LD blocks are set to zero. Bottom right panel shows a shrinkage-based estimate, where off-diagonal elements are shrunk towards zero.

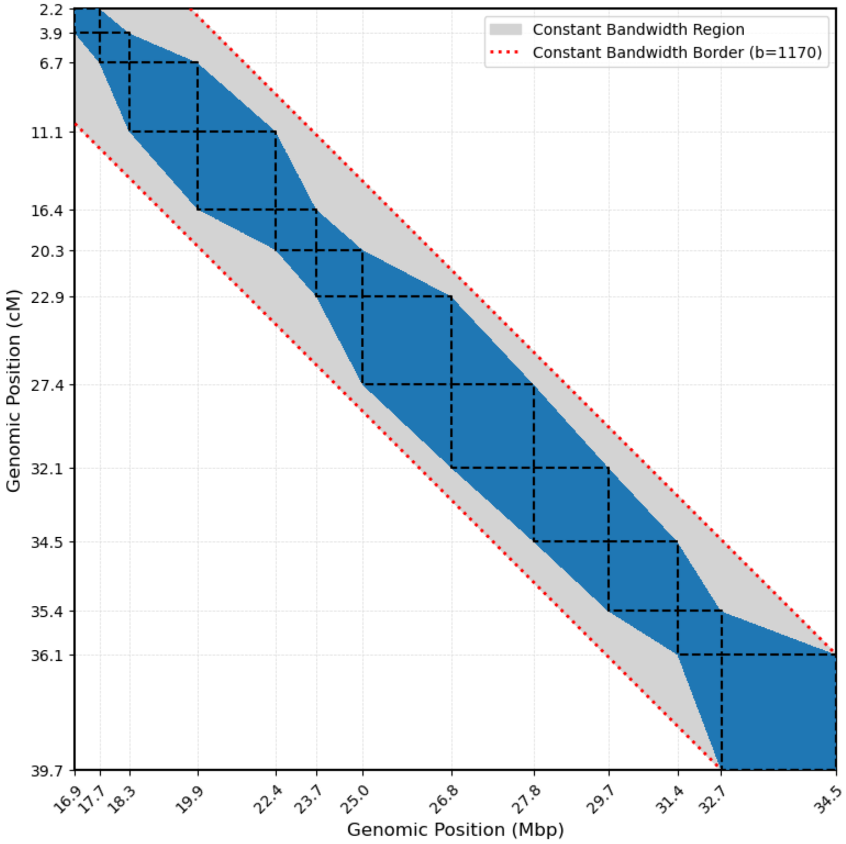


Figure S6: Visualization of the adaptive bandwidth mask on a region of Chromosome 22. The proposed mask (blue area) is constructed for a matrix based on a known block-diagonal structure. The original diagonal blocks are outlined with black dashed lines. The mask creates a continuous, non-uniform band by filling the space between these blocks. For comparison, the red dotted lines illustrate the constant bandwidth that is given by the largest block.

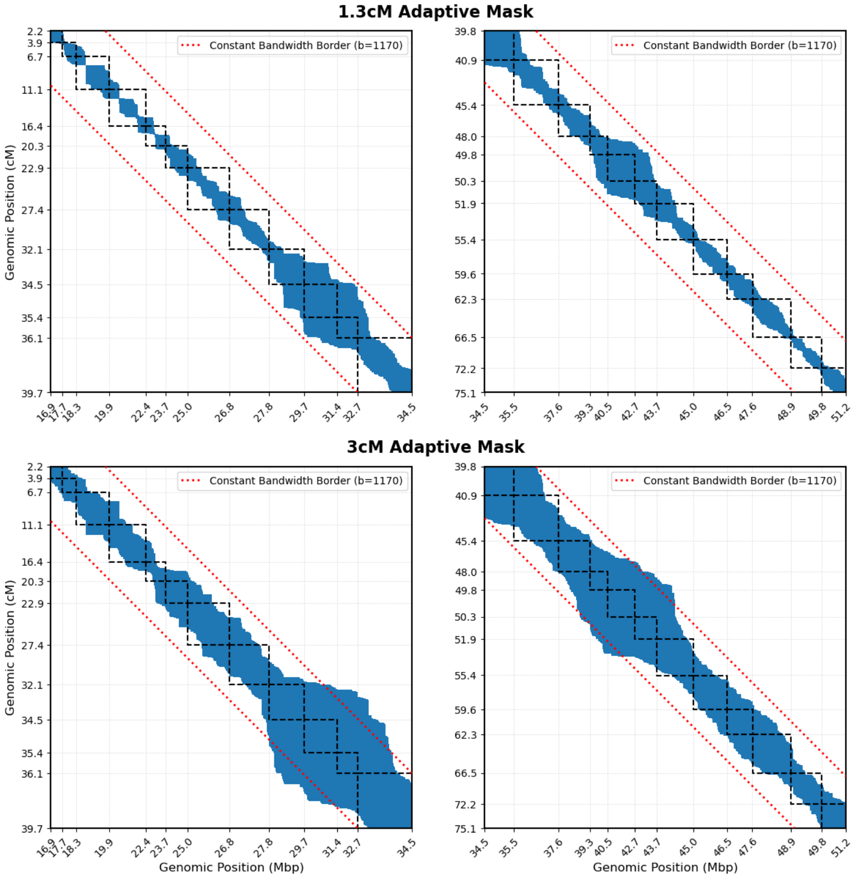


Figure S7: Comparison of adaptive sparsity masks derived from genetic distance of the complete Chromosome 22. The blue area in each panel represents the non-zero elements of a sparse mask where correlations are retained up to a fixed centiMorgan (cM) distance. For reference, we added dashed black squares showing the underlying block-diagonal structure and the red dotted lines indicating the borders of a constant bandwidth from Figure S6. (Top) A mask constructed with a 1.3 cM cutoff. This specific distance was chosen to match the total number of non-zero elements in the block-diagonal approximation (Table S1). (Bottom) A mask constructed with a more generous 3 cM cutoff, which retains more long-range correlations at the cost of being less sparse. The right plots are the continuation of the plots from the left.

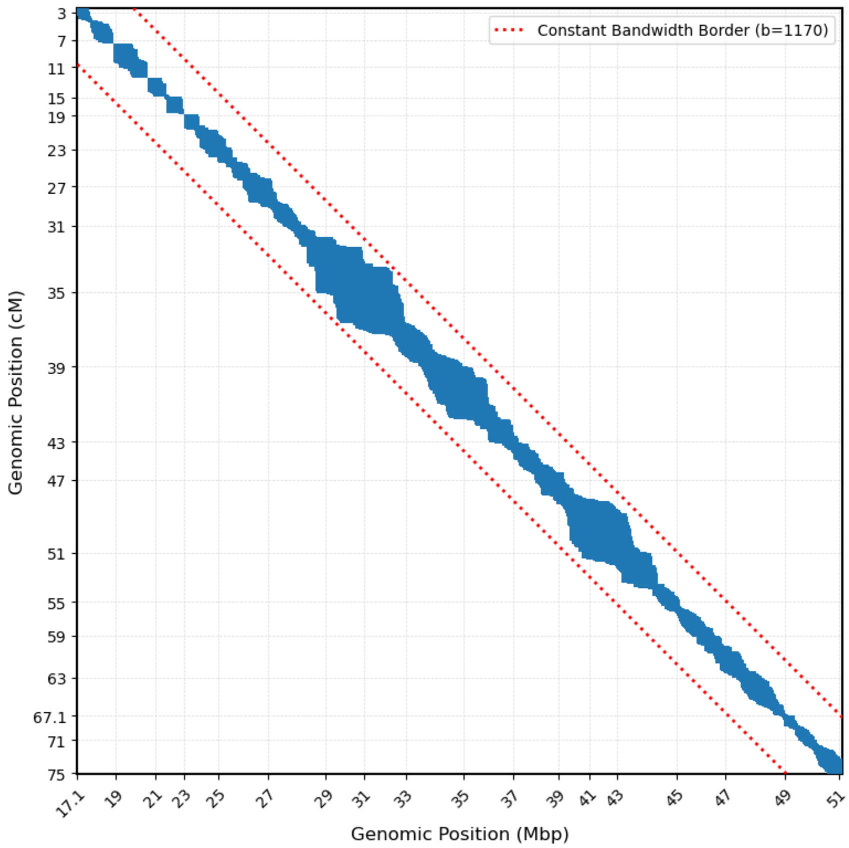


Figure S8: Relationship between distance measures among SNPs on Chromosome 22. The blue area shows the sparsity pattern of the 1.3 cM adaptive mask. Both x-axis and y-axis ticks represent variants chosen at uniform intervals according to a particular genomic distance. The x-axis uses ticks at approximately 2M base pairs (Mbp), while the y-axis uses ticks at approximately 4 cM. The grid cells shows the variation in recombination rates across the chromosome: regions where the rectangles are vertically stretched correspond to recombination "hotspots" (more cM per Mb), while vertically compressed regions correspond to "coldspots".

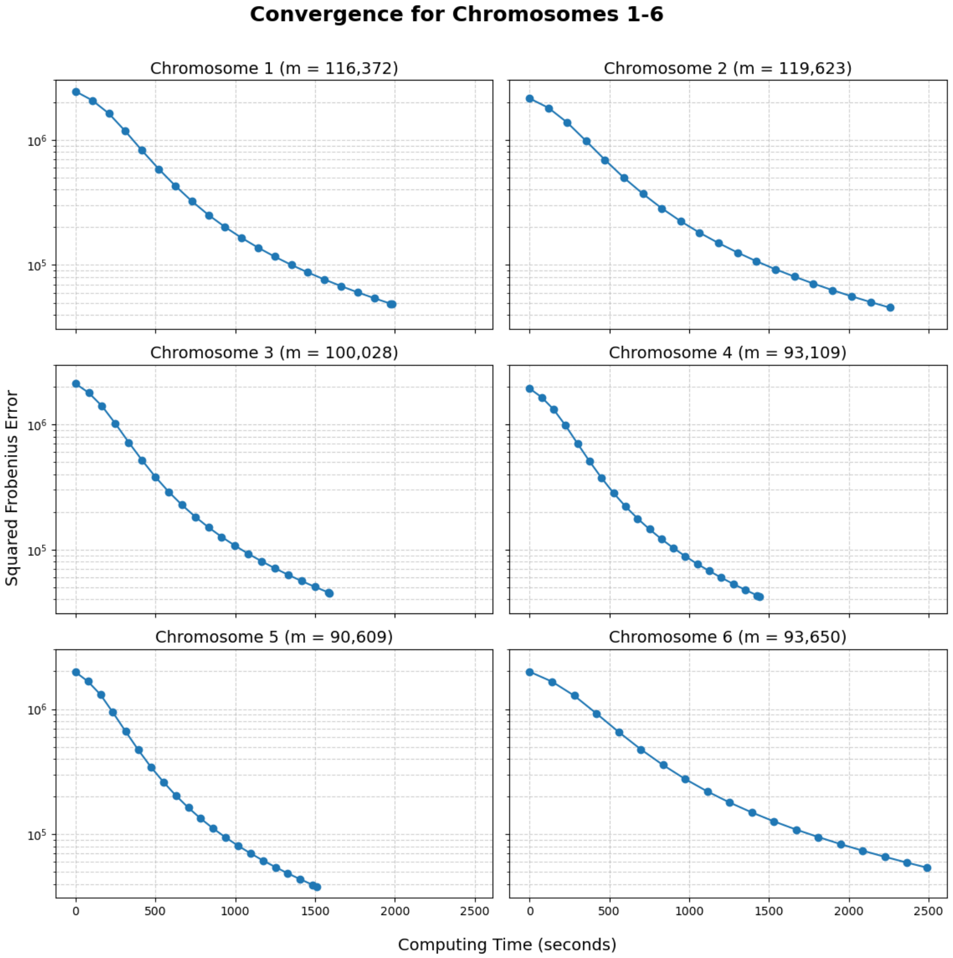


Figure S9: Convergence of the Cholesky-based algorithm on the six largest human autosomes. The plots show the value of the objective function given by the Squared Frobenius Error ($\|\mathbf{R}_b - \hat{\mathbf{R}}\|_F^2$) as a function of cumulative computing time in seconds, with the error axis presented on a logarithmic scale. On each panel, we can see the performance for Chromosomes 1 to 6, with the number of variants $m$ noted in the subtitle.

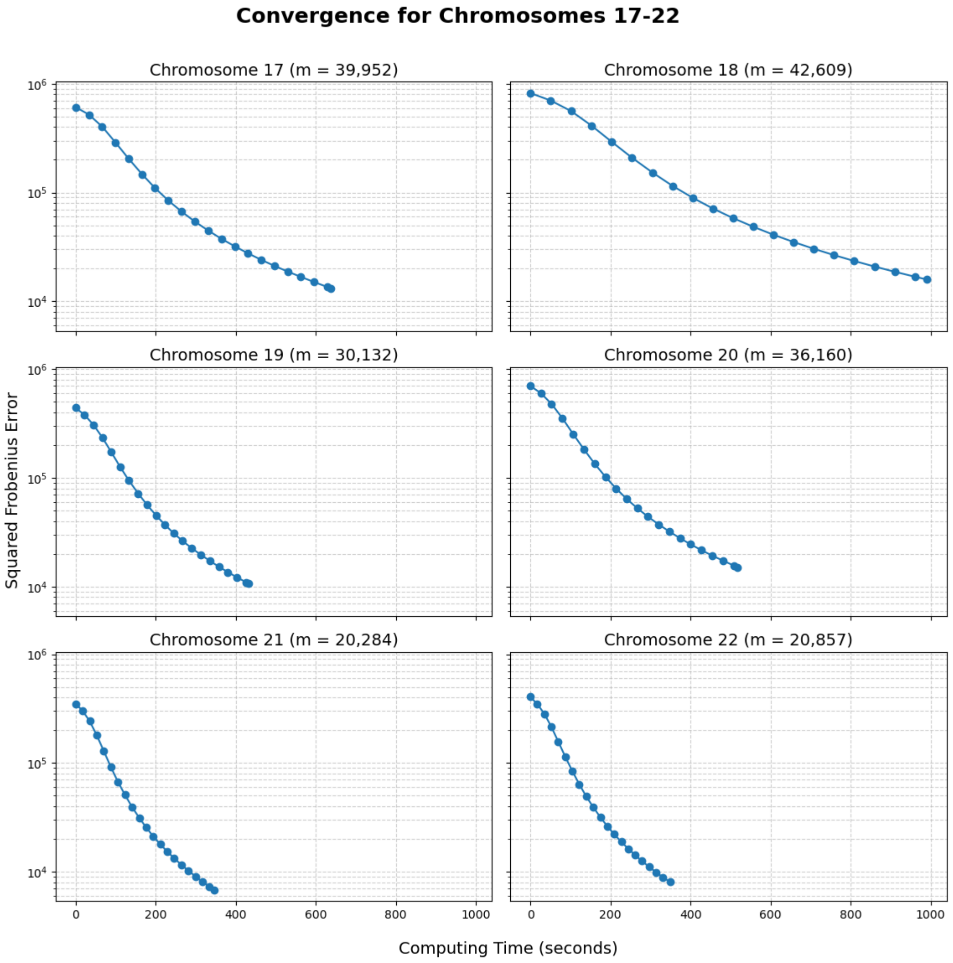


Figure S10: Convergence of the Cholesky-based algorithm on the six smallest human autosomes. The plots show the value of the objective function given by the Squared Frobenius Error ($\|\mathbf{R}_b - \hat{\mathbf{R}}\|_{\mathrm{F}}^2$) as a function of cumulative computing time in seconds, with the error axis presented on a logarithmic scale. On each panel, we can see the performance for Chromosomes 17 to 22, with the number of variants $m$ noted in the subtitle.

#### Supplementary Tables

| Approximation Method | Non-Zero Elements |
|---|---:|
| cholesky/windowed (1.3 cM) | 11,535,360 |
| cholesky/windowed (fixed band) | 35,940,788 |
| block-diagonal | 11,735,542 |
| shrinkage | 43,295,760 |

Table S1: Comparison of sparsity across different LD matrix approximation methods for Chromosome 22 ($m = 15,938$). The total number of elements in the dense matrix is $15,938^2 \approx 2.54 \times 10^8$.

| Chromosome | Variants (m) | Non-Zero Elements | Density (%) |
|---|---|---|---|
| 1 | $1.16 \times 10^5$ | $1.07 \times 10^8$ | 0.79 |
| 2 | $1.20 \times 10^5$ | $1.19 \times 10^8$ | 0.83 |
| 3 | $1.00 \times 10^5$ | $1.04 \times 10^8$ | 1.04 |
| 4 | $9.31 \times 10^4$ | $8.98 \times 10^7$ | 1.04 |
| 5 | $9.06 \times 10^4$ | $9.17 \times 10^7$ | 1.12 |
| 6 | $9.37 \times 10^4$ | $1.18 \times 10^8$ | 1.35 |
| 7 | $8.05 \times 10^4$ | $7.68 \times 10^7$ | 1.19 |
| 8 | $7.78 \times 10^4$ | $8.15 \times 10^7$ | 1.34 |
| 9 | $6.51 \times 10^4$ | $5.75 \times 10^7$ | 1.36 |
| 10 | $7.48 \times 10^4$ | $6.92 \times 10^7$ | 1.24 |
| 11 | $7.20 \times 10^4$ | $7.68 \times 10^7$ | 1.48 |
| 12 | $6.96 \times 10^4$ | $6.44 \times 10^7$ | 1.33 |
| 13 | $5.31 \times 10^4$ | $5.03 \times 10^7$ | 1.79 |
| 14 | $4.71 \times 10^4$ | $4.51 \times 10^7$ | 2.03 |
| 15 | $4.26 \times 10^4$ | $3.55 \times 10^7$ | 1.95 |
| 16 | $4.56 \times 10^4$ | $3.56 \times 10^7$ | 1.71 |
| 17 | $4.00 \times 10^4$ | $2.89 \times 10^7$ | 1.81 |
| 18 | $4.26 \times 10^4$ | $3.53 \times 10^7$ | 1.94 |
| 19 | $3.01 \times 10^4$ | $2.06 \times 10^7$ | 2.27 |
| 20 | $3.62 \times 10^4$ | $2.77 \times 10^7$ | 2.12 |
| 21 | $2.03 \times 10^4$ | $1.39 \times 10^7$ | 3.38 |
| 22 | $2.09 \times 10^4$ | $1.51 \times 10^7$ | 3.47 |

Table S2: Summary of matrix dimensions and sparsity for the 1 cM adaptive mask for each autosome. The table lists the total number of variants (m) for each chromosome and the corresponding number of non-zero elements in the resulting sparse approximation mask. The density column represents the percentage of non-zero elements relative to a dense m x m matrix.

| Chromosome | Computation Time (s) | | Peak RAM (GB) | |
|---|---|---|---|---|
| | Mean | Std. Dev. | Mean | Std. Dev. |
| 1 | 2005.6 | 12.30 | 28.43 | 0.89 |
| 2 | 2283.2 | 16.45 | 30.12 | 0.71 |
| 3 | 1604.6 | 14.38 | 25.55 | 0.35 |
| 4 | 1450.4 | 6.62 | 22.50 | 0.29 |
| 5 | 1518.2 | 6.69 | 22.94 | 0.17 |
| 6 | 2539.8 | 94.62 | 25.81 | 0.22 |
| 7 | 1420.0 | 6.04 | 21.35 | 0.38 |
| 8 | 1339.4 | 8.14 | 21.08 | 0.34 |
| 9 | 1404.8 | 2.17 | 17.69 | 0.05 |
| 10 | 1361.6 | 2.88 | 19.71 | 0.27 |
| 11 | 1580.8 | 6.61 | 20.56 | 0.28 |
| 12 | 1184.2 | 8.23 | 18.15 | 0.25 |
| 13 | 1027.0 | 4.58 | 15.13 | 0.16 |
| 14 | 841.0 | 6.44 | 12.54 | 0.20 |
| 15 | 899.2 | 2.77 | 10.79 | 0.15 |
| 16 | 982.8 | 1.30 | 10.98 | 0.11 |
| 17 | 637.4 | 1.82 | 9.96 | 0.18 |
| 18 | 989.0 | 2.55 | 10.91 | 0.21 |
| 19 | 429.6 | 1.82 | 7.72 | 0.20 |
| 20 | 517.4 | 1.52 | 9.61 | 0.19 |
| 21 | 344.2 | 1.79 | 7.21 | 0.22 |
| 22 | 348.4 | 1.14 | 7.06 | 0.19 |

Table S3: Computational benchmark for the Cholesky-based approximation across all 22 autosomes, showing mean and standard deviation over five replicate runs.

#### References


1. Golub, G. H. & Van Loan, C. F. *Matrix Computations - 4th Edition* (Johns Hopkins University Press, Philadelphia, PA, 2013).
2. George, A. & Liu, J. W. H. *Computer Solution of Large Sparse Positive Definite Systems* (Prentice Hall, 1981).
3. Harris, C. R. *et al.* Array programming with NumPy. *Nature* **585,** 357–362 (2020).
4. Joblib Development Team. *Joblib: running Python functions as pipeline jobs* 2020.
5. Kingma, D. P. & Ba, J. *Adam: A Method for Stochastic Optimization* 2017.
